# Microtissue geometry and cell-generated forces drive patterning of liver progenitor cell differentiation in 3D

**DOI:** 10.1101/2020.10.28.355875

**Authors:** Ian C. Berg, Erfan Mohagheghian, Krista Habing, Ning Wang, Gregory H. Underhill

## Abstract

Towards understanding the impact of mechanical signaling on progenitor cell differentiation in three dimensional (3D) microenvironments, we implemented a hydrogel based microwell platform to produce arrays of multicellular microtissues in constrained geometries, which cause distinct profiles of mechanical signals. We applied this to a model liver development system to investigate the impact of geometry and stress on early liver progenitor cell fate. We fabricated 3D liver progenitor cell microtissues of varied geometries, including cylinder and toroid, and used image segmentation to track individual single cell fate. We observed patterning of hepatocytic makers to the outer shell of the microtissues, except at the inner diameter surface of the toroids. Biliary markers were distributed throughout the interior regions and was increased in toroid tissues compared to cylinder tissues. Finite element models of predicted stress distributions demonstrated that cell-cell tension correlated with hepatocytic fate, while compression correlated with decreased hepatocytic and increased biliary fate. This combined approach integrating microfabrication, imaging and analysis, and mechanical modeling demonstrate of how microtissue geometry can drive patterning of mechanical stresses that regulate cell differentiation trajectories. It also can serve as a platform for the further investigation of signaling mechanisms in the liver and other systems.

## Introduction

The differentiation and morphogenesis of progenitor cells into functioning tissues is orchestrated through a diverse set of biochemical cues from neighboring cells and other elements of microenvironment. Mechanical force and mechanical signaling have been established as key to stem cell development and tissue behavior.^[1]^ Much of the work relating stem cell behavior to mechanical conditions, such as stiffness, geometry, and cell generated force, has been constrained to two-dimensional culture systems. Three-dimensional culture is more representative of in vivo conditions owing to increased cell-cell interactions and freedom for motility and reorganization, which are especially important in differentiation and morphogenesis. Compared with those on 2D, cells in 3D experience distinct mechanical stresses which influence their biological functions in embryonic development and tumor growth.^[2]^ As 3D culture platforms have become more widely utilized, those that more specifically address mechanical loading and signaling have also been developed.^[3]^

Since basic parameters, such as size, can affect cell and aggregate behavior^[4]^, careful engineering is required to investigate cell response to a 3D environment. Microwell based platforms have emerged as a method to create large numbers of replicate multicellular structures for use in drug screening, disease modeling, and stem cell culture. ^[5]^ Platforms involving ECM protein or hydrogel encapsulation have been used to demonstrate that geometry of an embedded 3D tissue can drive tube morphogenesis and cancer invasion through endogenous stress patterns.^[6]^ Other systems, utilizing collagen encapsulated tissues constrained by mechanically tuned pillars, have shown patterns of ECM protein organization in response to mechanical stress.^[7]^ Microwell systems, without an encapsulating matrix, have been suitable for engineering the mechanical condition as cell derived forces responding to the constraining geometries were sufficient to cause alignment of ECM proteins.^[8]^

Implementation of 3D systems with mechanical constraints to more tissue specific contexts, such as stem and progenitor cell development has been more limited. One system of interest is development of the liver. In the embryo, the hepatic diverticulum is comprised of bipotential progenitor cells, also referred to as hepatoblasts. These cells are capable of differentiation into either hepatocytes or biliary epithelial cells, also called cholangiocytes, and eventually establish the well described repeating lobule structure of the liver. The process of fate specification and eventual morphogenesis of liver tissue and bile ducts occurs in a spatially and temporally orchestrated process in the region immediately surrounding a portal vein, guided by a diverse set of cues. Hepatocytic fate is primarily associated with signaling through Wnt, HGF, and FGF.^[9]^ while biliary fate is linked to Notch and TGFβ activity.^[10]^

Using a model liver progenitor cell line, bipotential mouse embryonic liver (BMEL) cells^[11]^, we have previously demonstrated that attributes of the microenvironment, such as ECM protein composition and substrate stiffness, can further influence differentiation ^[12]^ Using this model cell line, we additionally characterized the relationship between increased cell to surface traction force with increased biliary differentiation at the periphery of circular, monolayer, multicellular islands, highlighting the importance of mechanical signaling in this process.^[13]^ Here, the traction force distribution was a consequence of the monolayer geometry, demonstrating a relationship between geometry, mechanical loading, and cell behavior in these liver progenitor cells, an attribute that has been exploited in other 2D stem cell culture systems.^[14]^ As in other organ systems, 3D culture of liver tissue is more physiologically relevant, and we have observed that differentiation of BMEL aggregates can also be affected by modifying the 3D environment.^[15]^ However, leveraging tissue geometry to alter mechanical signaling in 3D, has been more limited. Further, the relationships between geometry, mechanical signaling, and liver progenitor fate specification in 3D has not yet been characterized.

In this work, we implement an ECM-scaffold-free, hydrogel microwell based method to produced arrays of BMEL cell microtissues. The tissues have defined 3D geometries, including cylinders and toroids, with dimensions comparable to those of the liver lobule, which vary the predicted stress distributions. Using a single-cell image-segmentation-based approach, we characterize the 3D patterns of hepatocytic and biliary marker expression in this model liver progenitor cell line, when cultured in various tissue geometries. We also implement a finite element method model to predict the stress conditions of different regions of the tissues to investigate how tensile and compressive stress correlate with early progenitor cell specification. We also explore the dependence of these patterns on cell contraction. Our findings provide additional evidence that mechanical signaling plays a role in the 3D spatial patterning of progenitor cell fate specification in liver development. These findings highlight the importance of considering tissue geometry and resulting stress condition in 3D cell culture systems used to study stem cell behavior. Further, this combination of fabrication, imaging, and modeling, could serve as a platform to further investigate these relationships in this and other multicellular systems.

## Results

### Micromolded PEG substrates generate 3D microtissue with defined geometries

We implemented a microwell based approach to generate arrays of bipotential mouse embryonic liver (BMEL) cell three-dimensional tissues with defined geometries. For fabrication, a master wafer was prepared using photolithography and etching (Supplementary Fig S1A). Initial trials using single level PDMS molds required large volumes of high-density cell suspension, limiting the tissues that could be produced. This motivated incorporating a loading well similar to the seeding chamber provided by other available molds, which are produced by 3D printing rather than standard soft lithography.^[5b]^ To achieve a multilevel mold without additional lithography steps or other equipment, we first fabricated a 500 μm PDMS gasket sheet with an array of holes large enough to surround each sub-array of features. We passivated the gasket with a fluoro-silane. The gasket sheet was positioned over the wafer before addition of the PDMS (Fig 1A). After curing, the PDMS was removed from the wafer, and the gasket sheet was peeled from the rest of the PDMS producing the two-level mold (Supplementary Fig S1B).

**Figure 1.**
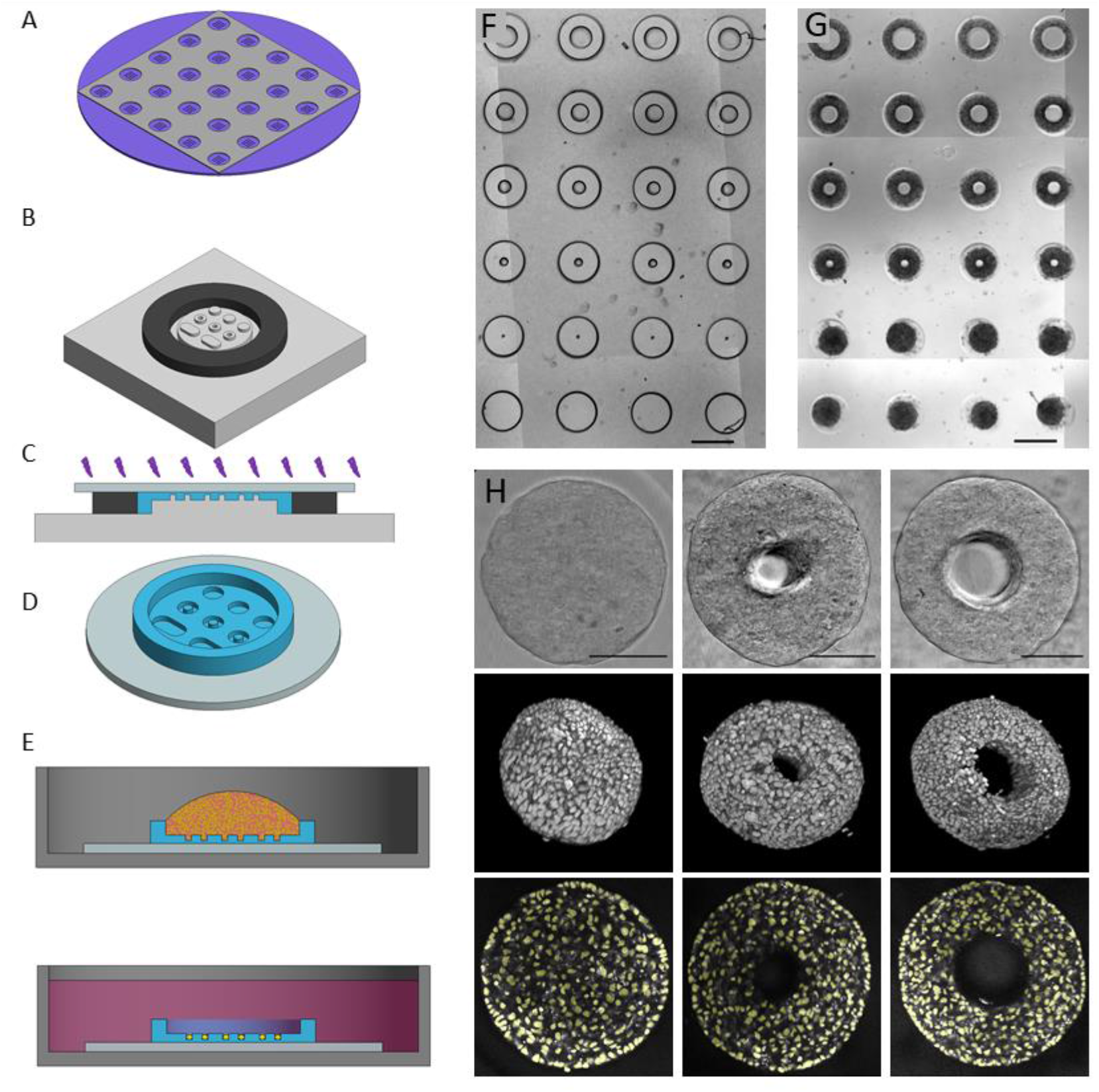
Micromolding-based process for microwell and tissue fabrication, and image analysis processes. A) A master wafer, prepared via photolithography and DRIE etching, is fit with a passivated PDMS gasket. PDMS is cured on the wafer to produce molds. B) A circular gasket is fit around the mold region of interest. C) The prepolymer solution is added to the mold, an activated coverslip is pressed on the droplet, and it is polymerized with UV light. D) Rendering of hydrogel substrate. E) For tissue seeding, a drop of cell suspension is added to the inset region of the substrate. Following centrifugation, excess cells are removed, and cells condense into aggregates. F) Brightfield image of a section of a PEG substrate with an array of microwells with equal cross-sectional area. G) Bright field image of an array of tissues. H). Top: Brightfield images of cylinder tissues (left), and toroid shaped tissues of varied inner diameter (middle and right). Center: 3D renderings of confocal images of nuclei distribution in tissues. Bottom: Nuclei image segmentation overlaid on central confocal image slices of tissues.

For substrate fabrication, a prepolymer solution consisting of 100 mg mL^-1^ 4-Arm-PEG-Acrylate (Laysan Bio) and 10 mg mL^-1^ photoinitiator (Irgacure 2959, BASF). A 1 mm thick PDMS gasket was positioned around the feature (Fig 1B, Supplementary Fig S1C). A droplet of the prepolymer solution was placed over the PDMS mold, a 12 mm acrylate functionalized coverslip was floated over the droplet, and the assembly was exposed to UV (Fig 1C). The coverslip was removed from the mold, resulting in a PEG substrate with 500 um side walls, with an inset array of microwells approximately 200 μm deep (Fig 1B, Supplementary Fig S1). This approach enables a wide variety of well geometries with high aspect ratio features, such as cylinder and toroid wells of varied inner and outer diameter (Figure 1D). In this example, the inner post diameter ranges from 50 to 250 um, and the outer diameter is adjusted such that all wells have the same cross-sectional area as a 400 μm diameter circle. This approach aims for tissues with approximately the same initial number of cells across the geometries. When measured optically, well dimensions closely reflected those drawn in CAD for the photomask (Supp Figure 1A, B).

For tissue fabrication, the substrates coverslips were placed into 24 well plate wells and sterilized with UV light. A droplet of cell suspension (~10 E6 cells mL^-1^) was placed in the loading well of each substrate. The plates were centrifuged driving cells to completely fill the microwells. Excess cells were removed with media washes (Fig 1E). Because the microwells were non-fouling and non-adhesive, cells only adhered to themselves, and aggregated into dense, 3D tissues constrained by the wells. In cylindrical wells, the cells aggregate and condense into a roughly cylindrical tissue. In the toroid microtissues, the cells aggregate and condense away from the outer walls, but around the central post, resulting in roughly donut shaped tissue (Fig 1G,H). Confocal images highlight the 3D structure of the microtissues (Fig 1H). After 72 hours in culture, the mean diameter of tissues in 400 μm diameter wells was 226 μm. the mean diameter of condensed tissues in toroid wells with 150 μm diameter posts, was 286 μm (Supplementary Fig S1G). Subsequent references to toroid and cylinder microtissues describe these two geometries unless otherwise noted. We used image segmentation^[14d]^ to identify and locate individual nuclei to enable single cell analysis in 3D (Fig 1H). We observed that despite the equal cross-sectional area, the mean final number of cells incorporated into cylinder tissues was higher than that in and toroid tissue, at 3498 and 2841 respectively (Supplementary Fig S1F).

### Hepatocytic and biliary marker patterning in cylindrical microtissues

In this work, we utilized bipotential mouse embryonic liver (BMEL) progenitor cells, which maintain bipotentiality when cultured at low density in growth conditions. These cells express increased markers of hepatocyte function when cultured in a differentiation media at high density and can be induced to express biliary markers in certain conditions.^[11–13]^ In previous work with BMEL cells, we correlated the expression of the hepatocytic marker, transcription factor HNF4a, with albumin expression, a marker of hepatocyte function. We have also correlated expression of cholangiocytic marker osteopontin (OPN) with the additional biliary markers SOX9 and Ggt1 .^[12a]^ Though during growth, BMEL cells can transitively express mixed markers, consistent with previous studies, in this microwell system we observed a large, correlated increase in HNF4a and albumin expression during 72 hours of culture, as measured by RT-qPCR. Concurrently, there is a gradual fall in OPN expression as the OPN expressed by unspecified cells is reduced even as some number of cells begins towards biliary fate, increasing their OPN expression (Supplementary Figure S2). Thus, in this study, we use HNF4a and OPN for markers of the early hepatocytic and cholangiocytic fate specification, which can be observed at 72 hours in these model liver progenitor cells.

To assess the distribution of cells with these markers within tissues, after 72 hours of culture, microtissues were fixed and immunostained for OPN and HNF4a. In tissues produced from 400 μm diameter cylindrical wells, cells expressing OPN were sparsely distributed throughout the tissue, while HNF4a expressing cells were almost exclusively found at the outer shell of the tissues (Figure 2A, B, Supplementary Figure S3). To quantify this behavior across microtissues, we implemented an image segmentation based single cell analysis pipeline to determine cell phenotype and location in a coordinate system normalized by the tissue radius and thickness (Supplementary Figure S4A). We mapped the percentage of HNF4a and OPN positive cells across a normalized radial and Z coordinate (RZ) space from replicate (N = 45) tissues, taking advantage of the rotational symmetry. This revealed consistent spatial patterning of HNF4a positive cells at the outer surface of the tissues (Figure 2D). Here, OPN positive cells were distributed across the cross section, with some exclusion from the outer region (Figure 2G).

**Figure 2.**
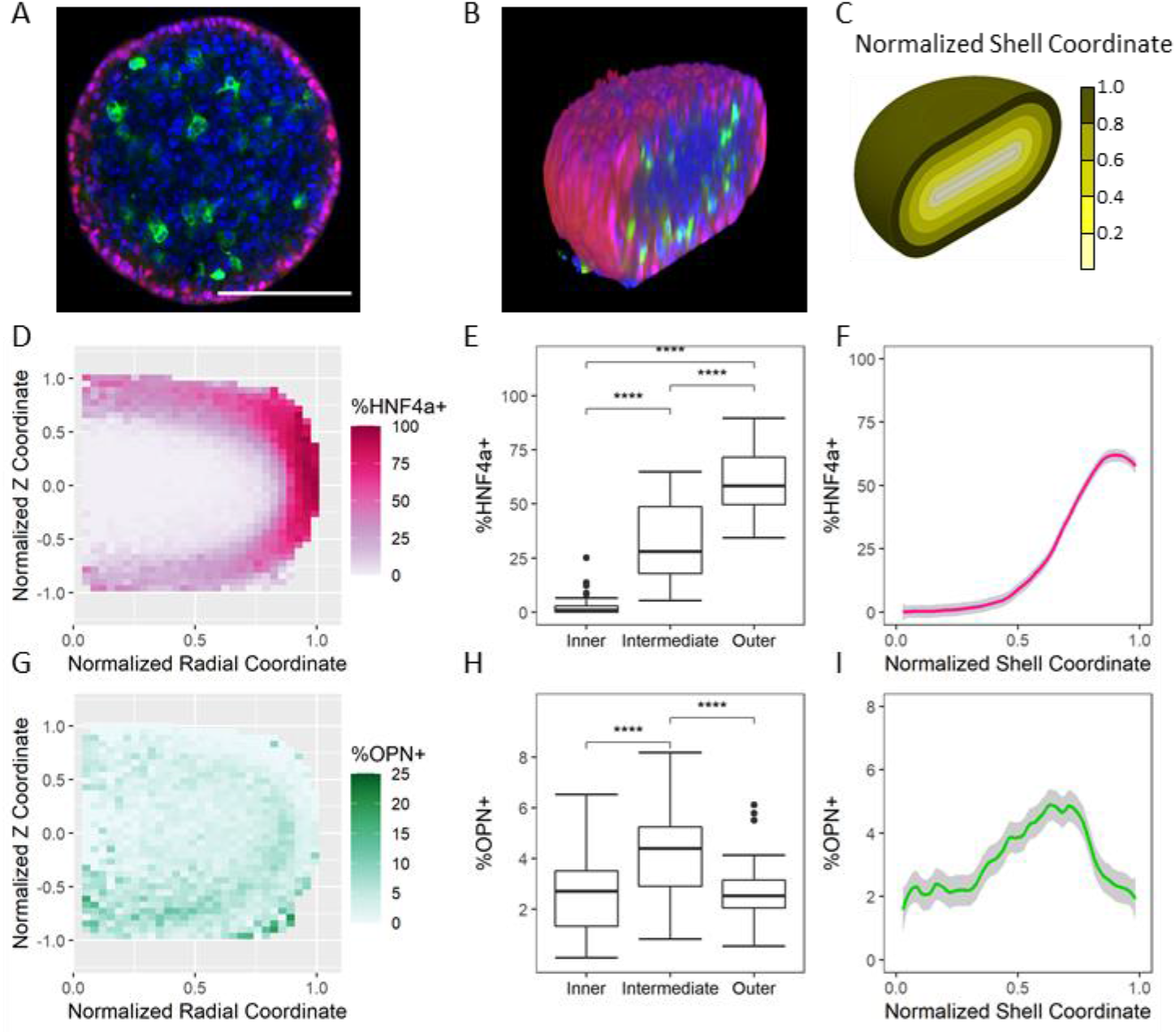
Spatial patterning of HNF4a and OPN expression in cylinder microtissues. A) XY confocal slice of a representative cylinder microtissue stained for hepatocytic marker HNF4a (red), biliary marker OPN (green), and with DAPI (blue). Scale bar 200 μm B) 3D rendering of confocal image stack, sliced across the approximate center XZ plane. C) Schematic view of the normalized shell coordinate. D) Percentage of HNF4a positive cells in RZ space from N=45 tissues. E) Percentage of HNF4a positive cells in the Inner, Intermediate, and Outer regions. F) Percentage of HNF4a positive cells versus normalized shell coordinate, mean ± 95% CI shown in gray. G) Percentage of OPN positive cells in RZ space. H) Percentage of OPN positive cells in the Inner, Intermediate, and Outer regions. I) Percentage of OPN positive cells versus normalized shell coordinate, mean ± 95% CI shown in gray. **** p <= 0.0001.

The condensed tissues adopt a final shape with a circular XY cross section, and a more ovular RZ cross section, which can be approximated as a rectangle with semicircular caps (Supplementary Figure S4B, C). Based on this approximation, we calculated a normalized shell coordinate to describe cell position with a single dimension (Figure 2C, Supplementary Figure S4C). Plotting percentage of HNF4a positive cells versus this coordinate further highlighted the outer surface patterning of HNF4a positive cells (Figure 2F). Plotting percentage of cells positive for OPN versus this coordinate revealed the decrease in OPN positive cells at the outer surface, and a slight elevation in OPN positive cells just inside of the outer region (Figure 2I).

Further, we sorted cells into an inner, intermediate, and outer region based on the shell coordinate (Supplementary Figure S4C) and calculated the percentage cells positive for each marker. We observed a statistically significant difference in percentage of HNF4a positive cells across all regions, with the outer region being the highest, and inner the lowest (Figure 2H). We observed a statistically significant increase in percentage of OPN positive cells in the intermediate region compared to the outer and inner regions.

### Hepatocytic and biliary marker patterning in toroid microtissues

Like the cylindrical tissues, in toroid tissues, produced with wells with a 150 μm center pillar, OPN positive cells were sparsely distributed across the 3D structure while HNF4a positive cells were primarily found at the outer surface. However, low levels of HNF4a positive cells were observed at the surface contacting the PEG pillar (Figure 3A, B, Supplementary Figure S5). Plotting percentage HN4a positive cells in RZ space from replicate (N = 40) microtissues demonstrates this behavior is consistent across tissues (Figure 3G). Plotting percentage of OPN positive cells in RZ space revealed reduced OPN positive cells in the outer shell, but not at the region contacting the pillar. As the PEG pillar is chemically inert to the cells, this region would be expected to behave as the rest of the outer surface, implicating mechanical interactions in altering the differentiation behavior.

**Figure 3.**
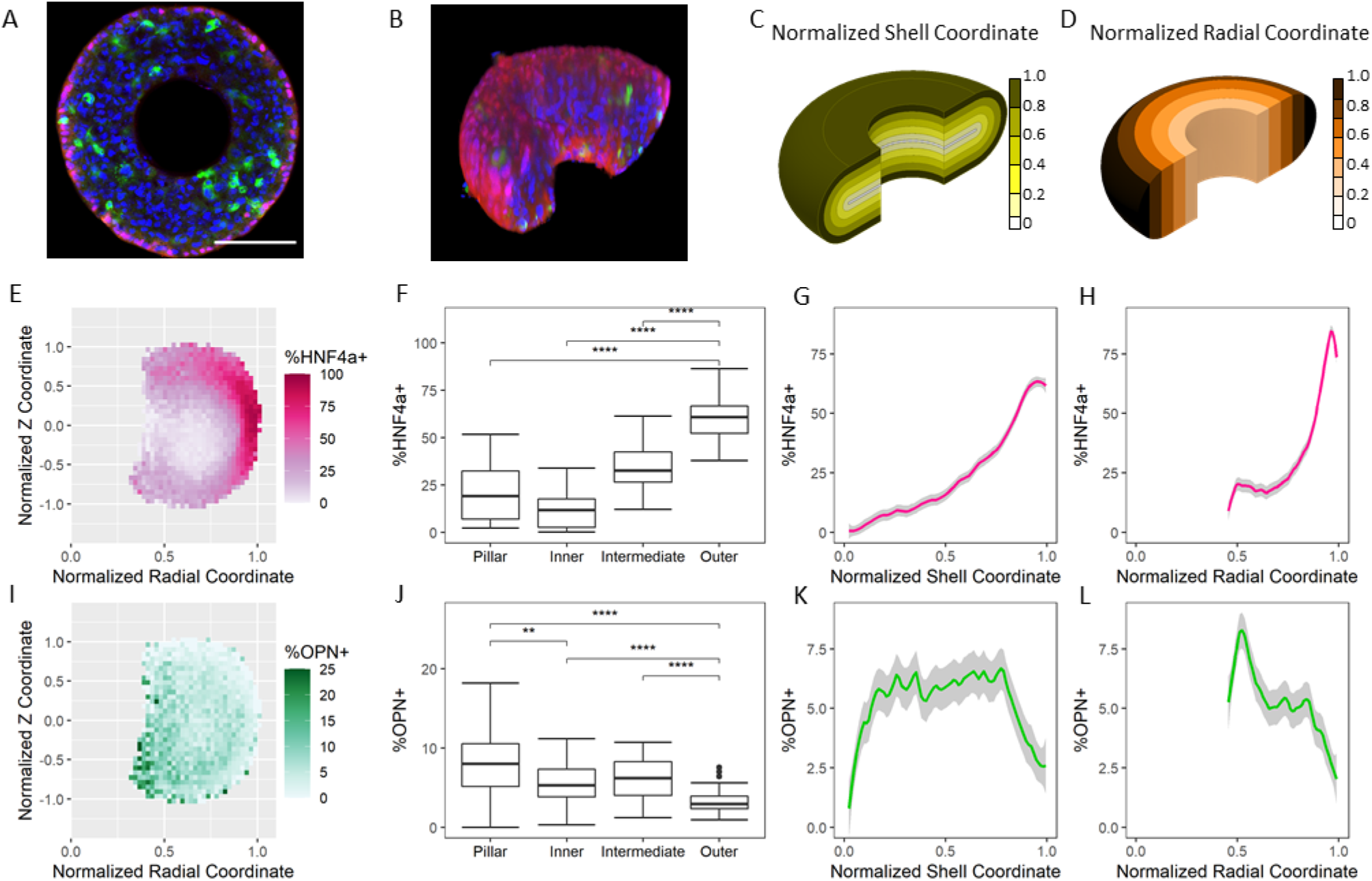
Spatial patterning of HNF4a and OPN expression in toroid microtissues. A) XY confocal slice of a representative toroid microtissue stained for hepatocytic marker HNF4a (red), biliary marker OPN (green), and with DAPI (blue). Scale bar 200 μm B) 3D rendering of confocal image stack, sliced across the approximate central XZ plane. C) Schematic view of the normalized shell coordinate. D) Schematic view of the normalized radial coordinate. E) Percentage of HNF4a positive cells in RZ space from N=40 tissues. F) Percentage of HNF4a positive cells in the Pillar, Inner, Intermediate, and Outer regions. G) Percentage of HNF4a positive cells versus normalized shell coordinate. H) Percentage of HNF4a positive cells versus normalized shell coordinate. Mean ± 95% CI shown in gray. I) Percentage of OPN positive cells in RZ space. J) Percentage of OPN positive cells in the Pillar, Inner, Intermediate, and Outer regions. K) Percentage of OPN positive cells versus normalized shell coordinate. L) Percentage of OPN positive cells versus normalized radial coordinate. Mean ± 95% CI shown in gray. ** p <= 0.01, **** p <= 0.0001.

As with cylinder microtissues, we use the plane of radial symmetry define a normalized shell coordinate (Figure 3C, Supplementary Figure S6A, B). Plotting marker expression frequency in this space emphasizes that HN4a positive cells are most found at the outer region, and the percentage of OPN positive cells drops in this region (Figure 3E, I). This coordinate does not indicate proximity to the pillar contacting surface. Thus, we also plotted percentage of cells positive for each marker against normalized radial coordinate (Figure 3D). This revealed that the percentage OPN positive cells increased near the pillar contacting surface, where R is approximately 0.4. Further, we used these two coordinates to sort cells into inner, intermediate, outer, and pillar regions (Supplementary Figure S6B). The percentage of cells positive for HNF4a was statistically significantly higher at the outer region compared to each other region, including the pillar. Percentage of OPN positive cells was lowest at the outer region. There was a small increase in percent OPN positive at the pillar compared to the inner region (Figure 3H).

### Toroid shape and EGF increase early biliary fate specification

Overall, untreated toroid microtissues had a statistically significant increase in percentage of OPN positive cells per tissue compared to cylinder tissues (Figure 4A). Treatment with EGF increased the percentage of OPN positive cells in both toroid and cylinder microtissues, while causing only a minor decrease in the percentage of HNF4a positive cells (Figure 4A, B). As with untreated tissues, EGF treated toroid tissues showed an increase in percent OPN positive cells compared to EGF treated cylinder tissues. EGF treatment did not disrupt the spatial pattern of HNF4a positive cells. In both the cylinder and toroid geometries, HNF4a cells were mostly restricted to the outer surface. In the EGF-treated toroid microtissues, HNF4a positive cells were restricted from the pillar contacting surface (Figure 4C, D).

**Figure 4.**
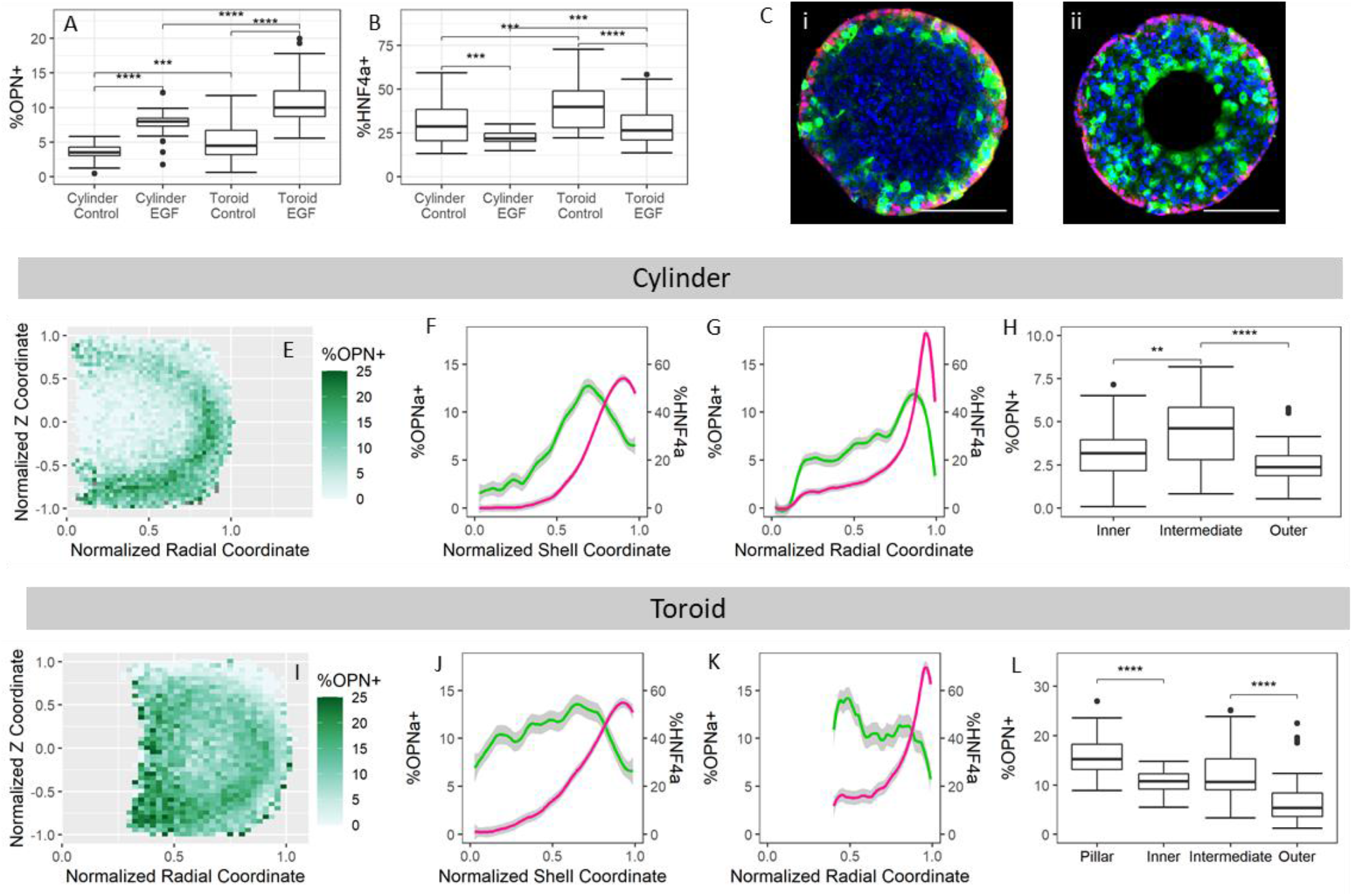
HNF4a and OPN expression in EGF treated BMEL microtissues. A) Percentage of OPN positive cells per microtissue by shape and treatment. Control is no treatment. B) Percentage of HNF4a positive cells per microtissue by shape and treatment. Control is no treatment. C) Confocal slice of a representative EGF treated cylinder (i) and toroid (ii) microtissue stained for HNF4a (red), OPN (green), and with DAPI (blue). Scale bar 100 μm. E) Percentage of OPN positive cells in RZ space from N=32 cylinder microtissues. F) Percentage of OPN (green) and HNF4a (magenta) positive cells versus normalized shell coordinate in EGF treated cylinder microtissues. G) Percentage of OPN (green) and HNF4a (magenta) positive cells versus normalized radial coordinate in EGF treated cylinder microtissues. Mean ± 95% CI shown in gray. H) Percentage of OPN positive cells in the Inner, Intermediate, and Outer regions of EGF treated cylinder microtissues. I) Percentage of OPN positive cells in RZ space from N=44 toroid microtissues. J) Percentage of OPN (green) and HNF4a (magenta) positive cells versus normalized shell coordinate in EGF treated toroid microtissues. K) Percentage of OPN (green) and HNF4a (magenta) positive cells versus normalized radial coordinate in EGF treated toroid microtissues. Mean ± 95% CI shown in gray. L) Percentage of OPN positive cells in the Pillar, Inner, Intermediate, and Outer regions of EGF treated toroid microtissues. ** p<= 0.01, ***p<=0.001, **** p <= 0.0001.

Along with amplifying overall expression of biliary marker OPN, treatment with EGF amplified the spatial pattern of OPN expression. Plotting OPN frequency in RZ space reveals that OPN frequency increases in the intermediate area, just inside the outer surface (Figure 4E). Using the normalize shell coordinate highlights that percent OPN positive is low at the outer shell, where HNF4a cell frequency is highest, peaks in the intermediate zone, and decreases moving towards the core (Figure 4F). The pattern also holds true when using the radial coordinate (Figure 4G) This trend is confirmed when comparing percent OPN positive in the inner, intermediate, and core regions, as percent OPN positive is highest in the intermediate zone (Figure 4H).

In toroid tissues, we see in the RZ space that percent OPN positive cells similarly increases in region just inside of the outer shell as well as near the pillar contacting region (Figure 4I). The increase in OPN cell frequency just inside of the outer shell is highlighted in the shell coordinate plot, where OPN cell frequency is low near shell coordinate of 1, where HNF4a cell frequency is highest. Percent OPN positive peaks near shell coordinate of 0.75. Plotting OPN cell frequency versus normalized radial coordinate shows the other peak in percent OPN positive, near the pillar the contacting region (R ≈ 0.4). These two peaks are confirmed when comparing the percent OPN positive in the pillar, inner, intermediate, and outer regions (Figure 4L).

### Increased E Cadherin expression colocalized with increased HNF4a positive cells

To further characterize the architecture of the tissues we stained tissues for actin and E-cadherin (ECAD) along with HNF4a. In cylinder microtissues, we observed an outer actin ring, visible in XY cross sections (Figure 5Ai, Supplementary Figure S6A). An actin ring similarly forms at the outer periphery in toroid microtissues, and at the inner, pillar contacting surface (Figure 5ii). ECAD was expressed at low levels throughout, but highest at the outer shell in both cylinder and toroid geometries (Figure 5A iii). In toroid microtissues, ECAD expression did not increase near the pillar contacting region (Figure 5A iv). Thus, the regions of increased ECAD expression correlated with the regions of increased HNF4a positive cells. Inspection of these regions with higher magnification confirmed these patterns and that the increased ECAD expression occurred at cell-cell junctions (Supplementary Figure S6A). To assess the consistency and 3D characteristics of this behavior, we averaged actin and ECAD stain intensity across the RZ plane of symmetry of the tissues and averaged this across multiple tissues. From this, we observe that the actin ring extends into an actin shell. Further, this confirmed the increased ECAD expression in the outer shell region, coincident with the region of increased HNF4a positive cells (Figure 5B).

**Figure 5.**
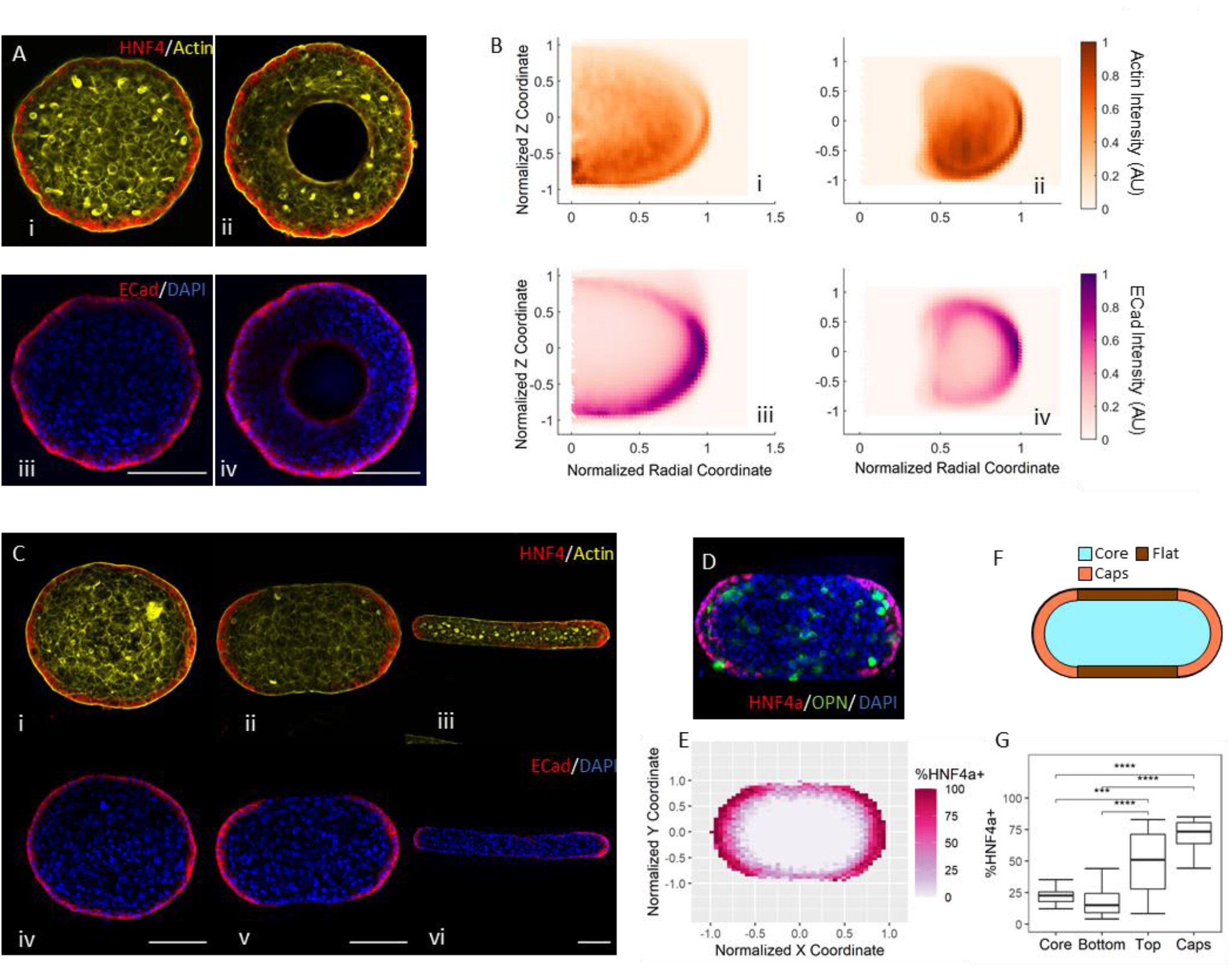
Patterning of actin, E-cadherin, and HN4a in 3D BMEL microtissues. A) Confocal image of a representative cylinder (i) and toroid (ii) microtissue stained for HNF4a (red) and actin (yellow), and with DAPI (blue), and a representative cylinder (iii) and toroid (iv) microtissue stained for ECAD (red) and with DAPI (blue). Scale bar 100 μm. B) Averaged actin intensity in RZ symmetry plane averaged from n = 8 (i) cylinder and (ii) toroid tissues in RZ space. Averaged ECAD staining intensity in RZ symmetry plane averaged from n = 8 (iii) cylinder and (iv) toroid microtissues. C) Confocal images of representative (i) 300 μm, (ii) 200 μm, and (iii) 100 μm wide oblong microtissues stained for HNF4a (red) and actin (yellow). Confocal images of representative (iv) 300 μm, (v) 200 μm, and (vi) 100 μm wide oblong microtissues stained for ECAD (red) and with DAPI (blue). Scale bar 100 μm. D) Confocal image of a representative 200 μm wide oblong microtissue stained for HNF4a (red), OPN (green), and with DAPI (blue). Scale bar 100 μm. E) Percentage of HNF4a positive cells in RZ space in the center 25% of the tissue height from N=18 200 μm wide oblong microtissues. F) Descriptive regions of oblong microtissues. G) Percentage of HNF4a positive cells in the core, flat, and regions of 200 μm tissues. ***p<=0.001, **** p <= 0.0001.

### HNF4a positive cells excluded from flat regions of oblong microtissues

To further explore the relationship between geometry and differentiation, we prepared “oblong” microtissues using wells with a racetrack like cross section, a rectangle with semicircular caps. Here, the wells were designed such that the width was set to 100, 200, or 300 μm, and the length is chosen such that the cross-sectional area is equal to that of a 400 μm diameter circle. After 72 hours of culture, tissues in the 300 μm wide oblong wells condense into slightly elongated, ovular cylinders. In these geometries, the spatial patterns of actin, ECAD, and HN4a expression was like those observed in cylinder microtissues (Figure 5C i,ii). Microtissues in the 200 and 100 μm wide oblong wells condense such that they are pressed against the side walls, producing microtissues with flat sides and rounded caps. In these tissues, the cortical actin ring is present around the entire periphery of the microtissue cross section. However, HNF4a positive cells were generally found at the round cap regions, and less so at the flat, wall contacting sides. Areas of increased ECAD expression also followed this pattern (Figure 5 iii-vi, Supplementary Figure S7A).

To confirm this phenomenon, we stained for and quantified HNF4a positive cells in replicate 400 μm oblong tissues (Figure 5D). Plotting percent HNF4a positive cells in normalized XY space for the central 25% of the tissues’ thickness showed that the oblong tissues consistently have higher levels HNF4a positive cells at the round caps than the flat sides, and HNF4a positive cells were infrequently found through the core (Figure 5E). Dividing the tissues into core, cap, and flat regions (Figure 5F), the caps have statistically significant higher levels of HNF4a positive cells than the flat and core regions (Figure 5G). Viewed in normalize XZ space, we observed that the top region of the tissues behaves similarly to the caps, while the bottom of the tissue behaved like the flat sides (Supplementary Figure S7C-F).

### Blebbistatin disrupts HNF4a expression patterning and OPN expression

To investigate the role of actin-myosin contractility on differentiation in 3D, we treated microtissues with blebbistatin to disrupt actin-myosin contraction. Overall, treatment with blebbistatin reduced OPN expressing cells in all geometries (Figure 6I), suggesting a role of cell contractility in biliary fate specification. In both cylinder and toroid microtissues, blebbistatin did not greatly affect overall levels of HNF4a expressing cells compared untreated or vehicle (DMSO) treated (Figure 6I) but did appear to disrupt the spatial patterning (Fig 6A, B). In the cylinder microtissues, there remained increase in HNF4a positive cells compared to the core, however this sorting was less pronounced in blebbistatin treated relative to DMSO treated microtissues (Figure 6C,D). Using the shell coordinate, we observed that the peak level of HN44a positive cells was reduced and spread further into the structure (Figure 6G). Blebbistatin was even more disruptive to the spatial patterning in toroid microtissues (Figure 6E,F). The peak level HNF4a positive cells were more uniformly distributed across the microtissue volume (Figure 6H). Treatment with ROCK inhibitor Y-27632 did not impact the overall frequency or spatial pattern of biliary or hepatocytic markers in cylinder or toroid microtissues (Supplementary Figure S8).

**Figure 6.**
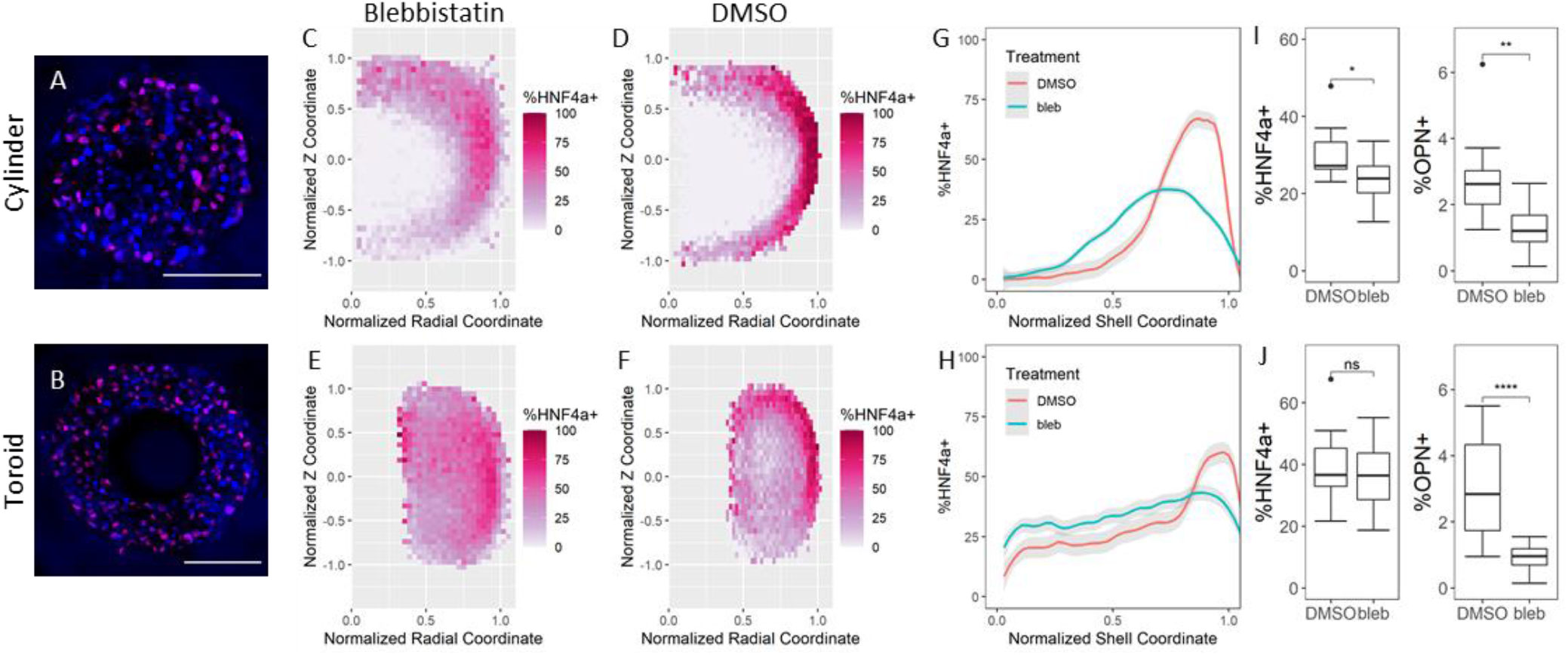
Disruption of HNF4a expression patterning with blebbistatin. Confocal image the central slice of a representative cylinder (A) and toroid (B) blebbistatin-treated microtissue stained for HNF4a (red) and with DAPI (blue). Scale bar 100 μm. C-F) %HNF4a positive heat maps for DMSO and blebbistatin treated cylinder and toroid microtissues. G,H) %HNF4a positive versus shell coordinate for DMSO and blebbistatin-treated cylinder and toroid microtissues. Mean ± 95% CI shown in gray. I,J) Overall %HNF4a and %OPN positive cells per microtissues in DMSO and blebbistatin-treated cylinders and toroids. * p<= 0.05, ** p<= 0.01, **** p <= 0.0001.

### Mechanical models suggest differing regions of compression and tension from cell derived forces

To understand the mechanical behavior of the microtissues, we implemented a 3D finite element method (FEM) based model, building off previously reported 2D and 3D tissue models (Supplementary Figure S9).^[14e, 16]^ Based on the actin and ECAD patterns observed, we modeled the microtissues as having a contractile shell, similar to that described in the aggregates produced and characterized by Lee et. al..^[16a]^ Based on the increased OPN positive cell frequency in the region just inside the outer shell, we modeled the intermediate region as stiffer than the core, also as suggested by Lee et. al. (Supplementary Figure 8A). The results of this model suggested that in the cylindrical microtissue, the outer shell region was primarily under tension, with positive first and second principal stresses, and third principal stress of approximately zero. The intermediate and core regions were entirely in compression, which the largest compressive stress in the intermediate region (Figure 7F, G, Supplementary Figure S10B). In the toroid microtissue model, the contractile shell region is also primarily under tension, and the intermediate and core regions were entirely in compression, with increased compression at the intermediate region. However, the pillar contacting region is experiencing compression at comparable levels to the intermediate zone (Figure 7H, I, Supplementary Figure S10B).

**Figure 7.**
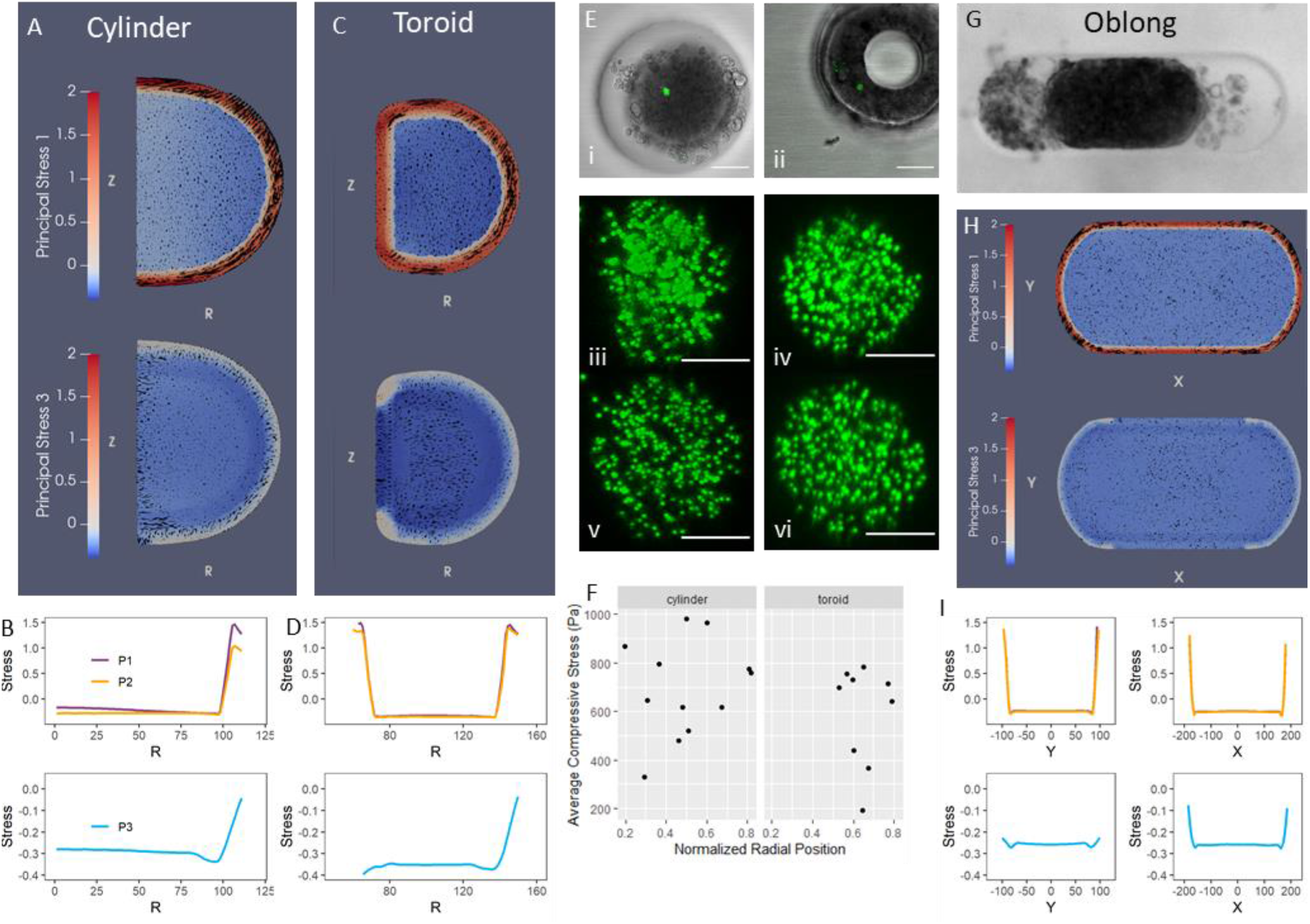
Modeling and measurement of stress in 3D BMEL microtissues. A) Simulated first and third principal stresses for an RZ slice of a representative cylinder microtissue. B) Simulated principal stresses at the line of Z = 0 from the above RZ slice (cylinder). C) Simulated first and third principal stresses for an RZ slice of a toroid microtissue. D) Simulated principal stresses at the line of Z = 0 from the above RZ slice (toroid). Stress values given in relative, arbitrary units. E) Representative cylinder (i) and toroid (ii) microtissues with an embedded microgel. Scale bar 100 um. Maximum intensity projections of the stressed (iii and iv) and relaxed (v and vi) microgels from the cylinder and toroid geometries shown above. Scale bar 10 um. F) Average compressive stress on microgels in cylinders and toroids plotted versus radial position. G) Brightfield image of an oblong microtissue. H) Simulated first and third principal stresses from the central XY slice of an oblong microtissue. I) Simulated principal stresses at the line of X = 0 (left) and Y = 0 (right) from the above XY slice (oblong).

To validate some of the results of this model, we imbedded alginate microgels into the microtissues, which contained fluorescent beads, to measure forces within the tissue. In these experiments, an image is acquired of the microgel under the stress of the tissue. The tissue is lysed with a detergent, and an image is collected of the relaxed (stress-free) microgel. The relative displacement of the fluorescent beads can be used to measure the cell forces of the microgel.^[2c]^ These microgels were successfully incorporated into both cylinder and toroid microtissues (Figure 7A). Visual inspection of the microgels before and after lysing revealed microgels frequently expanded upon relaxation, indicated they had been in compression. To assess this behavior, we measured the volume of the microgel before and after lysing. We then calculated the relative volume change and calculated the average compressive stress in each microgel. All measured microgels in both tissue geometries expand upon tissue lysing, indicating they were in compression. This confirmed the compression in the interiors of the tissues. The average compressive stress across replicate cylinder and toroid tissues was 695 ± 197 Pa and 590 ± 207 Pa respectively, however we did not observe a statistically significant difference in average compressive stress between toroid and cylinder tissues or as a function of radial position (Figure 7B). More dense distributions of microgels in toroid and cylinder tissues may be needed for studies in the future.

We next applied the FEM model to the oblong microtissues. After 72 hours of culture, the tissues continue to contact the long flat walls (Figure 7E). While this may be expected to occur during aggregation, under uniform contraction the tissue would be expected to uniformly shrink away from the walls. Under this model, there is no stress distribution (Supplementary Figure S10C). The continued pressing against the wall does occur when modeled as a contractile shell and passive core. When modeled in this way, without the walls of the well, we see this results in an outward “bowing” at the flat ends (Supplementary Figure S10D). In the oblong tissue model when the well wall is included, the contractile regions were primarily in tension. At the round caps, the third principal stress falls to near zero. The wall contacting regions were experiencing compression at similar levels to the intermediate and core regions (Figure 7J, K).

The results of our models suggest that geometry combined with the contractile shell drives regions of surface compression, which correlate to reduced hepatocytic differentiation and reduced ECAD expression. To explore this relationship further, we fabricated “double pillar” and “dumbbell” shaped microtissues (Figure 8). While similar to the oblong design, the model predicted the presence of the pillars cause the tissues to contract away from the walls, eliminating any compressed surfaces (Figure 8A). This behavior was confirmed experimentally (Figure 8B). Here we observed that the entire periphery contained increased HNF4a positive cells and increased ECAD expression, including the flat sides (Figure 8C,D). We also observed the cortical actin ring around the periphery (Figure 8H). The model predicts that the dumbbell microtissues contract such that only the center “bridge region” is compressed against the well wall (Figure 8D), which was confirmed experimentally (Figure 8F). Here increased HNF4a positive cells and ECAD expression was found around the periphery, except near this flat bridge region (Figure 8J,L). The cortical actin was also found at the periphery, with some decrease at these flat sections (Figure 8K).

**Figure 8.**
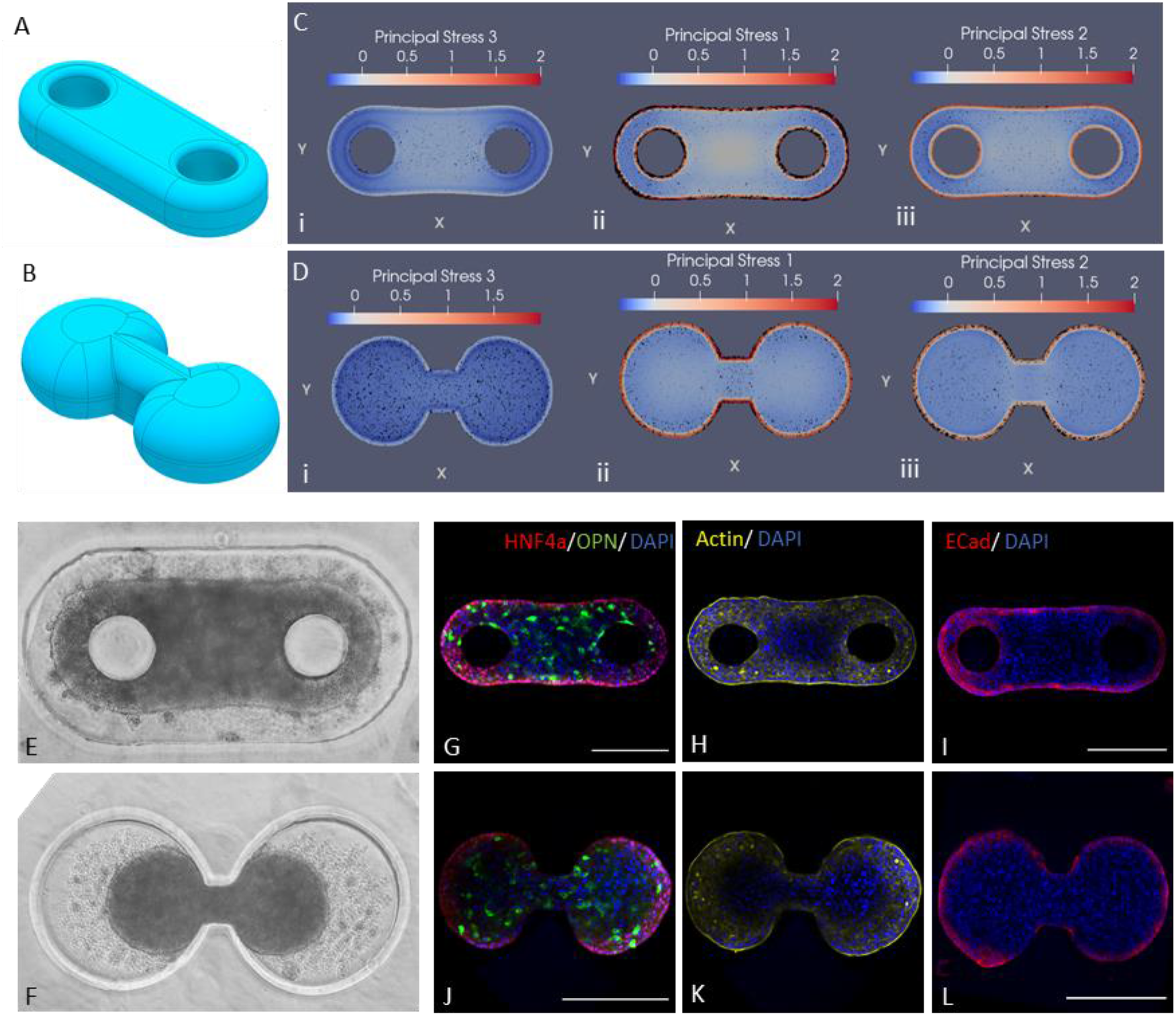
Behavior of double pillar and dumbbell microtissues. CAD model used for FEM simulation of the double pillar (A) and dog bone (B) tissues. Simulated principal stresses for the center XY slice of a (C) double pillar and (D) dumbbell geometries. Stresses are given in relative units. Bright field image of a (E) double pillar and (F) dumbbell microtissue. Confocal images the central slice of a representative (G-H) double pillar and (J-K) dumbbell microtissue stained for HNF4a (red), OPN (green), actin (yellow) and with DAPI (blue). Scale bar 200 μm. Confocal images the central slice of a representative (I) double pillar and (L) dumbbell microtissue stained for Ecad (red) and with DAPI (blue). Scale bar 200 μm.

## Discussion

### Method provides reproducible arrays of geometrically defined microtissues

The molded microwell-based microtissue fabrication method we implemented, when combined with the described imaging, analysis, and mechanical modeling pipelines, provided an effective platform for characterizing spatial patterns of differentiation in 3D as they related to geometry. The use of photolithography, etching, and soft lithography techniques produced consistent replicate microtissues with a variety of well-defined, high aspect ratio geometries. This included toroid geometries with 150 um inner diameter, which is comparable to the diameter of an embryonic mouse portal vein, providing for physiological relevance.^[17]^ It also enabled fabrication of cylinders, oblong shapes, dumbbells, and multi-pillar structures, which allowed us to impart varied compressive and tensile stress profiles.

Using the two step PDMS molding process outside of the cleanroom, we achieve a multilayer mold. The multilayer mold enables creation of a loading well, without the use of multiple lithography, mask alignment, and etching steps, or use of a 3D printer which while more versatile can be limited by resolution. This loading well permits smaller volumes for cell seeding and for immunostaining steps. The use of 4-Arm-PEG-Arcylate as the hydrogel base provides a suitably, non-fouling material with limited swelling to allow resolving small features. It is also readily modified with conjugation of biomolecules via Michael type addition, or inclusion of acylated molecules during polymerization, to make the wells adhesive or otherwise reactive, if desired for future applications. Though not considered in this study, the stiffness of the hydrogel can be modulated by varying molecular weight and monomer density of the PEG solution. This is relevant in the toroid and oblong wells, where the microtissue exerts force against the well and may experience an effect of the substrate stiffness.

Due to the modularity of the design and ease of implementation, these methods can be easily adapted to other 3D cell culture applications. Use of embryonic stem cells, induced pluripotent stem cells, or other primary progenitor cells in this system, possibly with an encapsulating matrix such as Matrigel, could be used to produce standardized embryoid bodies or organoids. Controlling 3D geometry could prove a powerful tool in guiding morphogenesis. The platform could also be used to generate standardized 3D tumor models of controlled geometry and characterize or screen for any impact of geometry on drug efficacy.

### 3D Liver progenitor cell microtissues form contractile shells with increased early hepatocytic fate specification

In the 3D liver progenitor cell microtissues, we observed a distinct spatial pattern of cells positive for HNF4a, a marker in this system of early hepatocytic fate specification, at the outer shell of cylinder tissues, with significantly reduced HNF4a positive cells through the core. This was accompanied by an increase in e-cadherin expression, and a layer of increased actin expression. This pattern was consistent with a model of mechanical polarization at the boundary causing an effective tissue surface tension.^[18]^ Under this model, the outer shell develops a higher degree of contractility owing to the cortical actin layer, causing the cells to have high tension resolved across cell-cell interfaces.^[19]^ Development of a tissue surface tension is especially expected in embryonic tissues, possibly due to osmolarity differences between interstitial fluid and cell culture media.^[20]^. Recent work by Lee et al. directly measured this phenomena in similarly formed tissues.^[16a]^ These results suggest that cell-cell tension correlates with hepatocytic differentiation. This is supported by our observation that hepatocytic cell patterning at the outer surface was reduced by treatment with blebbistatin. This is consistent with our previously reported observations in 2D constrained islands, where the central region showed the highest hepatocytic differentiation,^[13]^ where the highest cell-cell tension is generated. ^[21]^

### Reduced hepatocytic fate specification in compressive regions

Similar to the cylinders, in the toroid microtissues, we observed high levels of cells expressing the early hepatocytic differentiation marker HNF4a at the tissue-media interface region of the outer shell, and low levels throughout the core. Additionally, HNF4a expression was low at the region contracted against the center PEG pillar. Since the PEG hydrogel used is non-adhesive and non-fouling, biochemically this interface would be expected to behave like the tissue-media interface. Thus, we would expect to observe similar mechanical polarization and high hepatocytic differentiation. The difference between these regions demonstrates an impact geometry and mechanical loading. The pattern of increased ECAD expression was similarly diminished at the surface contracting against the pillar. The cortical actin shell in these tissues was somewhat reduced in this region.

Using microgel force sensors, we observed that the interiors of both tissue types were in compression. This supports the model of a contractile shell. It also suggested that the regions low in hepatocyte markers were those experiencing compression. We developed a finite element model in attempt to further understand this behavior. We had initially chosen the toroid geometry to mimic the portal vein region, and the prediction from previous models that a contracting toroid around a pillar yields nonuniform stress.^[8]^ Based on the patterning of phenotype, e-cadherin, and actin, and the microgel results, we modeled the tissues with a contractile shell and passive core similar to models described previously.^[16a]^ Under these assumptions, the outer shell is experiencing tension across what would be cell-cell interfaces, and the core is entirely in near uniform compression. In the toroid model, the entire shell is in cell-cell tension, and the core is in near uniform compression. However, at the pillar contacting boundary, the tissue experiences a significant compressive stress normal to the boundary. This suggests that compression, whether experienced across the core or against the well feature, decreases the likelihood of early hepatocytic specification. Treatment with blebbistatin allows hepatocytic differentiation to permeate further into the microtissue structure, possibly due to reduced cell force derived compression, supporting this hypothesis.

Our observations of progenitor differentiation within the oblong microtissues provide further support for the FEM model assumptions and relation to phenotype. The oblong microtissues, once aggregated, continue to contact the walls at the flat region throughout culture. This final geometry would be predicted during condensation under a tissue surface tension model, as the tissues approach an energetically favorable spherical configuration, constrained by the well. However, if contraction occurs following condensation, uniform contraction would cause the tissues to contract away from the walls, which has been observed in tissues of other cell types.^[5b]^ The behavior of being “pressed” against the flat walls only occurs given a contractile shell and passive core, which predicts that compressive stresses occur at the flat sides, normal to the boundary. These compressed regions correlate to the regions of reduced HNF4a and e-cadherin expression at the boundary. In the double pillar tissues, where the condensation leads to contraction away from the peripheral walls, there is no predicted compression and no region of excluded HNF4a expression. Though the mechanism relating compression to reduced hepatocytic differentiation remains unknown, recent work has demonstrated that compression can alter transcriptional behavior and cause depolymerization of actin.^[22]^ Here the tensile contractile shell and the resolved compressive regions follow from mechanical polarization at the boundary. This is often associated with tissue-media boundaries in 3D culture, however mechanical polarization at tissue-tissue boundaries has been observed in vivo, causing local increases in cortical tension and taking part in cell sorting.^[23]^ While in this system the PEG boundaries provide a non-adhesive service unlike what would be encountered in vivo, compressive forces can be achieved within adhesive matrices in some geometries, and thus may arise in vivo.^[16b]^

It is also important to note that in this system lack of curvature is closely tied to compressive regions. There is growing literature that cells and tissues respond to surface curvature in the microenvironment.^[24]^ The flat compressive regions in oblong microtissues have zero curvature, while the pillar contacting region has non-zero median curvature but zero Gaussian curvature.^[16a]^ There is evidence that cells respond differently to these curvature types.^[25]^ The double pillar microtissue results suggest that flat regions are still pushed towards hepatocytic fate in the absence of compression. However, when free from the it may be possible for sufficient curvature can form. Additionally, in 2D monolayers, hepatocytic differentiation occurs despite the flat plane geometry. However, it is possible that the response to curvature or relationship between curvature, stress, and phenotype are sensitive to other variables such as the adhesion molecule (i.e. integrin vs. cadherin) or stiffness.

### Increased early biliary fate specification with EGF, toroid geometry

The ability to produce consistent, replicate 3D microtissues and spatially analyze single cell behavior enable characterizing the more subtle effect of geometry on biliary differentiation. Biliary differentiation, as identified by OPN expression was found throughout the inner region of the structures, regions expected to be under compression. Overall, toroid microtissues had a higher level of biliary differentiation. Our FEM model predicts that the compression across the core regions is higher in toroid compared to cylinder geometries, possibly explaining this difference. In both geometries, we observed increased biliary differentiation in the intermediate region just inside the outer shell. In the toroid tissues, biliary differentiation also increased near the surface contracting against the pillar. Within the 3D microtissues, these regions are predicted to be under the most compressive stress. Treatment with blebbistatin, which reduces compression induced by cell forces, reduces biliary differentiation, supporting the link between biliary phenotype and compression. We directly measured compression within the tissues using alginate microgels. The imaged microgels throughout the interior of the tissues expanded upon tissue lysing, indicating they had been compressed by cell forces. This measurement technique does have several limitations. We did not track the relative vertical position of the microgel in the tissue, which the immunostaining data indicates is important. Further, the positions of the analyzed microgels are random, and based on bot uptake and ability to be imaged, the latter which is likely biased towards the bottom of the tissue where less penetration is required for imaging. Future work using a more optimized implementation of this or other emerging techniques to measure cell generated forces in 3D aggregates^[2c, 16a, 26]^ may provide a more quantified, precise stress map of the tissue to validate or refine our mechanical model. Nevertheless, the elastic round microgels reveal that the cylinder and toroid microtissues exert an average compressive stress of 200 to 1000 Pa, which is on the same order of magnitude as the shear storage modulus of the normal rat liver tissues^[27]^, suggesting that our microtissues could mimic at least partial behavior of the living tissues.

Previous experiments utilizing 2D circular islands demonstrated a link between increased cell-substrate traction force and increased biliary fate. Here, blebbistatin treatment decreased overall biliary differentiation and eliminated the pattern.^[13]^ Recent work has shown that cell to substrate tensile forces can impose vertical compressive forces on the nucleus, which can alter chromatin condensation, transcription factor activity, and gene expression, providing a possible link between our 2D and 3D observations.^[22, 28]^ Studies using embryonic stem cells progenitors found that compressive stress can induce upregulation of SOX9, a transcription factor associated with biliary differentiation.^[29]^ In bone remodeling and homeostasis, OPN is upregulated in response to compressive loading and is key to transduction of mechanical signals.^[30]^ Though outside of a liver context, these findings support the mechanically sensitive nature of some biliary differentiation pathways. Other recent work has demonstrated that nuclear envelope deformation allows cells to sense compressive forces, a possibility in these dense, crowded tissues.^[31]^ These data demonstrate that compression may be an additional tool to control or pattern differentiation or eventual morphogenesis in liver development or other stem cell and regenerative systems. This can be implemented in through physical compression or osmotic stress.^[29, 32]^

In all geometries tested, there were significant proportions of cells that, at the 72-hour time point, did not express either Hnf4a or OPN at high enough levels to be seen with the immunostaining used. Our hypothesis is that these cells maintain their bipotentiality and display distinct differentiation kinetics. Future assessment using additional markers or culture of disaggregated tissues may provide further insight on the cells in these regions. Treatment with EGF increased biliary differentiation in all microtissue geometries, suggesting these cells maintain their differentiation potential but may require additional factors or treatments to proceed at higher efficiency. EGF also amplified the spatial patterning aiding in characterizing these patterns. EGF has been previously demonstrated to play a role in both hepatocyte and biliary epithelium formation.^[9a]^ Our results add to growing evidence that EGF plays a role in biliary fate specification.^[33]^ EGF has been used in various progenitor and human pluripotent stem cell lines to generate cholangiocytes in vitro.^[34]^ More specifically, studies have reported EGF and EGFR induction of Notch signaling in the context of fate specification and morphogenesis.^[35]^ Further, Notch signaling is known to be mechanically sensitive ^[36]^, thus there may be synergies between EGFR/Notch signaling with mechanical stress in 3D. This is one example of mechanisms which can be explored in this system. Modulation of other pathways through drug treatment, siRNA, or genetic editing, would permit exploring their relationships to force and geometry. Reliance on immunohistochemical staining can partially limit the refinement of differentiation kinetics. In future efforts, the integration of relevant fluorescent reporters with this platform would enable live cell imaging to track differentiation patterning over time and other dynamic processes.

## Conclusion

We have implemented a microwell-based method to fabricate and analyze the behavior of geometrically defined 3D liver progenitor microtissues. We modeled the mechanical behavior with a finite element method model. We observed in this system that cell derived forces create regions of varied compressive and tensile stresses, which combined with the tissue geometry further drove spatial patterns of markers of early hepatocytic and biliary differentiation in liver progenitor cells. This relationship between mechanics and differentiation has not been described in a 3D system, and these findings provide insight into how geometric features may have a role in orchestrating liver development through mechanical signaling. Understanding these mechanisms could be leveraged in addressing developmental disorders and liver cancers, or in developing approaches to manufacture artificial livers from stem cells. Further, this integrated fabrication and analysis platform provides a framework to investigate the interactions between differentiation and mechanical loading in other stem and progenitor cell systems.

## Materials and Methods

### Wafer and PDMS mold fabrication

The master wafer was prepared using soft lithography techniques.^[37]^ A photomask was drawn in AutoCad and printed on to transparent plastic (CAD/Art services) which contained arrays of the desired geometries. A thin layer of negative photo resist (KMPR) was spin coated on to a clean 4” silicon wafer, and flood exposed to UV through the photomask. Following development, the exposed features were etched with deep reactive ion etching (STS Pegasus, SPTS Technologies) to a depth of approximately 200 μm.

For PDMS mold fabrication, a sheet of PDMS was first molded between two glass slides using a 500 μm spacer. 6 mm diameter holes, large enough to surround the feature arrays, were punched into the sheet. It was then treated with oxygen plasma and incubated with Trichloro(1H,1H,2H,2H-perfluorooctyl)silane vapor under vacuum for at least one hour. The sheet was positioned over the master wafer such that the holes aligned with the feature arrays to be molded. PDMS was poured over the wafer and sheet, fully degassed, and cured at 70 C for at least 3 hours. After curing, the 500 μm sheet was carefully peeled from the rest of the cured PDMS leaving 500 um raised platforms, which create to the loading well, on top of which the feature arrays protrude an additional ~200 um. The two-step PDMS molding technique enables creation of multiple mold levels without requiring additional photolithography or etching steps. Additionally, a PDMS sheet was molded between glass slides using a 1 mm spacer. This sheet was cut into individual gasket pieces, each with a 7 mm hole.

### Preparation of micro molded PEG substrates

Polyethylene glycol (PEG) hydrogels substrates were prepared by adapting previous protocols.^[5a-c, 38]^ 12 mm circular coverslips were immersed in 0.1 N NaOH for 1 hour, rinsed with DiH2O and placed on a hot plate at 110°C until dry. The NaOH treated coverslips were activated by immersion in 2% (v/v) 3-(trimethoxysilyl)propyl methacrylate in ethanol and placed on a shaker for 30 min. The activated coverslips were immersed in ethanol on the shaker for 5 min and again dried on a hot plate at 110°C. A 111.11 mg mL^-1^ 10kDa 4-Arm PEG-acylate (Laysan Bio) prepolymer solution was prepared in 1x PBS. This solution was mixed with Irgacure 2959 (BASF) solution (100 mg mL^-1^ in methanol) at a volumetric ratio of 9:1 (prepolymer to Irgacure) to achieve a final PEG concentration of 100 mg mL^-1^. The solution was then degassed under vacuum.

For PEG molding, the 1 mm gasket was positioned around the feature array to be used, leaving approximately 1 mm between the raised platform and the gasket. A 50 μL droplet of prepolymer solution was deposited on the mold. A pipette tip was used to spread the prepolymer solution across the mold and knock bubbles from the mold features. The droplet was covered with an acrylate coverslip, which was lightly pressed against the gasket. The assembly was exposed to UV light using a Spot UV Curing System (OmniCure S1500, Excelitas Technologies) with a 320-390 nm Filter and adjustable collimating adapter, at an intensity of approximately 50 mJ cm^-2^ for 30-60 seconds. The coverslip was then carefully removed from the mold and immersed in PBS in a 24 well plate. Prior to use in culture, the substrates were sterilized by immersion in PBS supplemented with 1% (v/v) pen/strep under UVC for 30 minutes.

### Cell culture and tissue formation

The experiments utilized BMEL 9A1 cells of passages between 28 and 34. BMEL cells were cultured as previously described.^[11a]^ For expansion, cells were thawed onto tissue culture plastic flasks coated with collagen I (0.5 mg ml^-1^) and incubated at 37°C and 5% CO2. For subculturing, flasks were treated with trypsin-EDTA (0.25% [v/v]) for ≤10 min to detach cells and replated on collagen I coated flasks. Growth media, for expansion, consisted of RPMI 1640 + GlutaMAX (Life Technologies, 61870-127) with fetal bovine serum (10% [v/v], FBS), penicillin/streptomycin (1% [v/v], P/S), human recombinant insulin (10 μg ml^-1^, Life Technologies, 12585-014), IGF-2 (30 ng ml^-1^, PeproTech, 100-12), and EGF (50 ng ml^-1^, PeproTech, AF-100-15).

For tissue formation, cells were collected and resuspended to ~10 E6 cells mL^-1^ in differentiation media. A 50μL drop of cell suspension was added to the recessed loading well of each substrate. The 24 well plate was then centrifuged to drive cells into the wells. Excessed cells were removed by repeated media rinses. Tissues were cultured in differentiation media at 37°C and 5% CO2 for 72 hours with a media change after the initial 24 hours. Differentiation media consisted of Advanced RPMI 1640 (Life Technologies, 12633-012) with FBS (2% [v/v]), P/S (0.5% [v/v]), L-glutamine (1% [v/v]), and minimum non-essential amino acids (1% [v/v], Life Technologies, 11140-050).

### Growth factor and drug treatments

All growth factors and drugs were prepared and reconstituted according to the instructions of the manufacturers: EGF (PeproTech, AF-100-15) 50 ng ml^-1^ in PBS with 1% w/v BSA; (-)-blebbistatin (Cayman Chemical, 13013), 1 mg ml^-1^ in dimethyl sulfoxide (DMSO); Y- 27632 (Enzo Life Sciences, 270-333-M005), 5 mg ml^-1^ in deionized water (DiH2O). Drugs were added to differentiation media at the following concentrations: (-)-blebbistatin, 25 μM; Y-27632, 15 μM. In experiments with addition of soluble treatments, these were added immediately after the excess cell media rinses. Media and treatments were refreshed after the initial 24 hours.

### Immunostaining

Cells were treated with brefeldin A (10 μg ml^-1^, R&D Systems, 1231/5 in differentiation media) for 2 h prior to fixation by immersion in paraformaldehyde (4% [v/v] in 1 × PBS) for 30 min at room temperature. Fixed samples were permeabilized by immersion in Triton X-100 (0.5% [v/v] in 1 × PBS) for 1 hour at room temperature. Samples were incubated in blocking buffer (5% [v/v] donkey serum in 1 × PBS) for 1 hour at room temperature. Primary antibody solutions were prepared by diluting one or more of the following antibodies in blocking buffer: mouse anti-HNF4a (1/200 from stock, Abcam ab41898), goat anti-OPN (1/60 from stock, R&D Systems, AF808), goat anti-Ecadherin (1/50 from stock, R&D Systems AF748), Acti-stain 488 phalloidin (7/1000 from stock, Cytoskeleton PHDG1-A). A 50 μL droplet of primary antibody solution was added to the recessed loading well of each substrate. Samples were incubated overnight at room temperature on a shaker. Samples were rinsed via 3 x 15-minute washes in PBS on a shaker. Secondary antibody solutions were prepared by diluting one or more of the following secondary antibodies in blocking buffer: DyLight 550-conjugated donkey anti-mouse IgG (1/50 from stock, Abcam, ab98767) and DyLight 488-conjugated donkey anti-goat IgG (1/50 from stock, Abcam, ab96935). A 50 μL droplet of secondary antibody solution was added to the recessed loading well of each substrate. Samples were incubated overnight at room temperature on a shaker. Samples were rinsed via 2 x 15-minute washes in PBS on a shaker. Samples were incubated in DAPI solution (Invitrogen D1306) for 1 hour, and briefly rinsed in PBS.

A droplet of liquid mountant (ProLong™ Diamond Antifade, Initrogen) was added to each substrate, and the coverslips were mounted onto standard microscope slides. Mounted samples were cured for at least 24 hours at room temperature. Once cured, samples were sealed with clear nail polish.

### 3D Tissue Imaging, Segmentation, and analysis

The fluorescently stained samples were imaged using a laser scanning confocal microscope (Leica TCS SP8). 3D image stacks were segmented into individual cells using an adaptation of the MINS (modular interactive nuclear segmentation) MATLAB/C++-based segmentation platform^[14d]^ on the nuclear image (DAPI) channel. Utilizing the Bio-Formats software tools^[39]^, the microscope software generated .lif files were directly loaded into the software to sequentially segment all image stacks from each tissue array, and automatically read in metadata such as pixel size.

Semi-automated post processing and analysis was completed using a series of MATLAB scripts. First, an edge detection-based process was used to identify the outer tissue boundary in each image slice to mask out cells and debris not incorporated into the tissue. Due to nature of the microwells, tissues can move and rotate in all three dimensions, thus rotation and alignment is required to compare spatial behavior across replicate tissues. For this, a best fit ellipse was fit to the XZ, YZ, and XY projections of the masked, segmented nuclei data. The orientations of the best fit ellipses were used to rotate the nuclei centroids such that major axes from the XZ and YZ projections were aligned with the horizontal (X) and vertical axis (Y) respectively, the major axis of the XY projection was aligned to the horizontal (X) axis. The nuclei were also translated such that tissue center is at the origin. The major and minor axis of the XY projection were used to normalize the X and Y coordinates, and the height of the tissue was used to normalize the z coordinates. This normalized coordinate system was used in subsequent analysis to enable spatial comparison across replicates independent of slight differences in tissue size after cell aggregation.

For OPN and HNF4a expression analysis, the average intensity of the relevant stains was calculated in each of the segmented nuclei, along with intensity of the surrounding 3D area. The difference between the intensities and the surrounding background were used for analysis to account for intensity difference due to imaging penetration across the thickness of the tissue. A two component gaussian mixture model was applied to these relative intensities within each tissue to sort tissues into positive or negative for each stain.

For phenotype heat map generation, cells from replicate tissues were binned in 2D according to the relevant normalized coordinates, and the percentage of cells within each bin positive for the strain was calculated. For 1D plots, normalized shell and radial coordinates were calculated as described in Supplementary Figure S4. For region-based box plots, regions were separated based on normalized radial and shell coordinates as shown in Supplementary Figure S4.

For e-cadherin and actin heat map generation, the corresponding images were rotated and centered using the segmented nuclei projection best fit ellipses as described earlier. Images were down sampled by a factor of 4, the XYZ coordinates for each remaining voxel were calculated, and coordinates were normalized using the best fit ellipses. Normalized XY coordinates were converted to a normalized radial coordinate. Voxels were binned into a triangle mesh, and the average intensity in each mesh unit was calculated. This was repeated across multiple tissues using the same mesh to generate an average heat map.

### RNA Isolation, RT-qPCR Analysis

For measurement of liver markers via RT-qPCR, microwell substrates were prepared containing arrays of only cylinder or only toroid wells, and cells were seeded into the wells to form tissues as described above. At the specified time point, the substrate was transferred to a well containing trizol solution (Life Technologies 15596-026), to collect RNA from the tissues within that substrate. For RNA isolation, samples were treated with DNAse (New England Biolabs, M0303S) at 37°C for 30 min and further purified using an RNeasy Mini Kit (Qiagen, 74104) following manufacturer’s instructions. Supermix iScript cDNA synthesis kit (BioRad, 1708841) was utilized to produce cDNA from isolated RNA. This was mixed with SsoAdvanced Universal SYBR Green Supermix (Bio-Rad, 1725264) and primer pairs at a final concentration of 100 nM (primer pair sequences listed in Supplementary Table 1). Thermal cycling and amplification curves measurement was completed using a CFX Connect Real-Time PCR Detection System (Bio-Rad). mRNA expression was calculated relative to Hprt1 and control samples as indicated using Bio-Rad CFX Manager 3.1 software.

### Finite element modeling

3D Tissue geometries were drawn in Autodesk inventor with dimensions that approximate measured final tissue dimensions. Separate solid bodies were generated for each material type in the tissue model. These geometries were exported as STEP or IGES files. The finite element mesh generator software Gmsh was used to generate tetrahedral mesh files for each CAD geometry.^[40]^ The finite element model was produced using the FEBio software suite.^[41]^ The PreView software was used to import the geometry, set boundary conditions, and specify material properties. Tissues were modeled as a union of bodies with each body being assigned a material.^[16a]^ The core and intermediate regions were modeled as Neo-Hookean solids. The free outer shell and pillar contacting shell regions were modeled as solid mixtures of a Neo-Hookean material and material with a prescribed isotropic active contraction. Due to the actual stiffness of the tissues being unknown, non-dimensional stiffness and prescribed contraction values were used to analyze the relative stress patterns arising from parameter choices. Intermediate, outer shell, and pillar-contacting shell regions were assigned a Young’s modulus of 1. The core Young’s modulus was set to 0.5. The outer shell and pillar contacting regions’ prescribed stress was set to 5. Because the PEG substrate is much stiffer than the tissue, the microwell boundaries and pillars were either modeled as rigid material with a sliding contact, or a boundary condition preventing displacement normal to the surface. The simulation was run using the FeBio solver as a steady-state static single time point using default settings. Simulation results were visualized in the PostView program, which was also used to export results as a VTK file. Paraview was used to calculate principal stresses, visualize data, and generate figures.^[42]^

### Microgel force sensor experiments

For microgel force experiments, the micowell substrates were fabricated on a glass bottom 35 mm dish (Cell E&G, GBD00003-200). Alginate microgels with fluorescent particles were fabricated as previously described with 0.5% w/v alginate.^[2c]^ The droplet suspension was mixed with the cell suspension just before seeding the microwells. Alginate is not stable in RPMI medium. Therefore, in microgel experiments, differentiation media was altered to use Advanced DMEM (Gibco 12491-023) instead of advanced RPMI. Images were acquired using a laser scanning confocal microscope (Leica TCS SP8). Microgels were first located and a low zoom image was collected to be able to mark the XY location of the particle within the tissue. Images of the stressed elastic round microgels were collected, and the tissues were treated with 2.5% Triton X-100 detergent to obtain stress-free conditions. Images of the particles were collected ever 5-10 minutes for approximately 1 hour until the stress-free condition was achieved. ImageJ was used to measure the diameter and centroid location of the tissues and the location of the microgel. 3D confocal images were deconvolved (AutoQuant X3), and the volume was measured using MATLAB scripts. Average compressive stress was calculated based on the Poisson’s ratio of 0.4 and the bulk modulus of 3012 Pa, from interpolation of previous alginate bead stiffness measurements.^[2c]^

### Statistical analysis

Experiments consisted of three or more biological replicates with multiple tissues per experiment. For percent positive heat map generation, all cells from all replicate tissues in the shown condition were binned in a 50 by 50 RZ space, percent of cells positive for each marker was calculated at each bin location and plotted. For region-based calculations, the percent positive cells in each region for each individual tissue was calculated and plotted. For line plots, the cells in individual tissues were binned in the relevant coordinate space, and a percent positive for the marker was calculated for each bin from each individual tissue. Line plots represent the mean percent positive for each bin position across replicate tissues with the 95% CI ribbons in gray. Line and box blots were generated using the ggplots2 package for R.^[43]^ For hypothesis testing, Welch’s two-sample t-test was performed using the base t.test function in R. p<0.05 was considered significant. P-values are indicated in the figures.

## Disclosures

The authors indicate no potential conflicts of interest.

## Acknowledgements

This work was supported by the National Science Foundation #1636175 to GU and by NIH GM 072744 to NW. Research reported in this publication was supported by the National Institute of Biomedical Imaging and Bioengineering of the National Institutes of Health under Award Number T32EB019944. The content is solely the responsibility of the authors and does not necessarily represent the official views of the National Institutes of Health. We gratefully acknowledge Hélène Strick-Marchand and Mary C. Weiss (Institut Pasteur) for the bipotential mouse embryonic liver (BMEL) cells. We acknowledge the Imaging Technology Group at the Beckman Institute for Advanced Science and Technology for help and advice with confocal microscopy.

## CRediT authorship contribution statement

**Ian C. Berg**: Conceptualization, Methodology, Software, Validation, Formal analysis, Investigation, Data Curation, Writing – Original Draft, Writing – Review & Editing, Visualization, Project administration

**Erfan Mohagheghian**: Methodology, Software, Resources

**Krista Habing**: Investigation

**Ning Wang**: Methodology, Resources, Writing – Review & Editing, Supervision, Funding acquisition

**Gregory H. Underhill**: Conceptualization, Methodology, Validation, Resources, Writing – Review & Editing, Supervision, Funding acquisition

## Data availability

Data available on request from the authors.

---

**Supplementary Figure S1.**
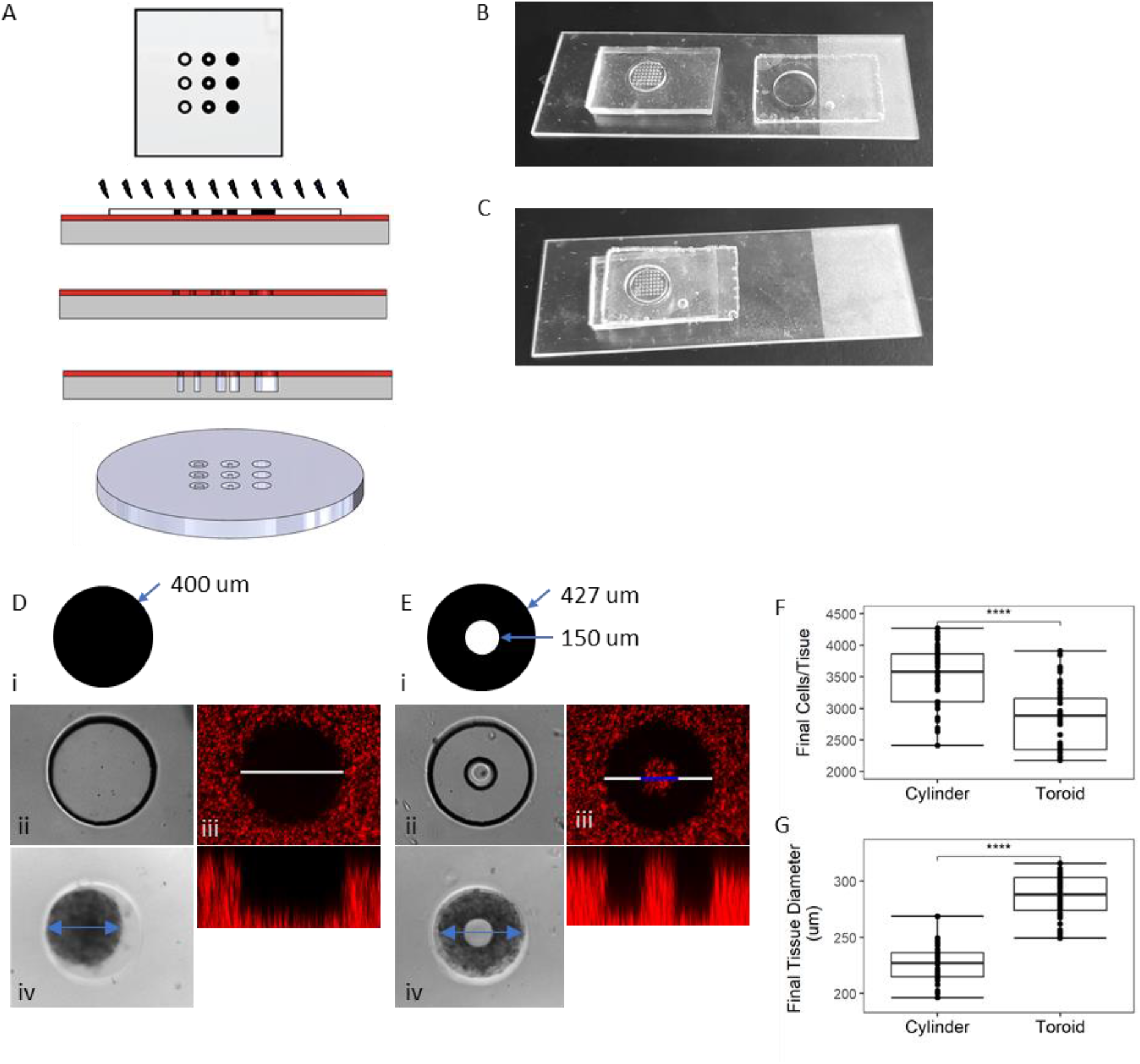
Fabrication process and characterization of hydrogel substrates and microtissues. A) Schematic of master wafer preparation by photolithography and etching. B) Multilevel PDMS mold of one feature array. C) PDMS mold with 1 mm gasket. D) Characterization of cylinder well. i) Feature design used in mask. ii) Bright field image of cylinder well. iii) XY (top) and XZ (bottom) confocal images of cylinder well with fluorescent beads mixed in hydrogel. White bar indicates 400 μm. iv) Brightfield image of cylinder tissue. Blue line indicates tissue diameter. E) Characterization of toroid well. i) Feature design used in mask. ii) Brightfield image of toroid well. iii) XY (top) and XZ (bottom) confocal images of toroid well with fluorescent beads mixed in hydrogel. White and blue bars indicates 427 and 150 μm. iv) Brightfield image of toroid tissue. F) Number of cells counted from segmentation per tissue after 72 hours of culture. G) Final outer diameter of tissues after 72 hours of culture. **** p <= 0.0001.

**Supplementary Figure S2.**
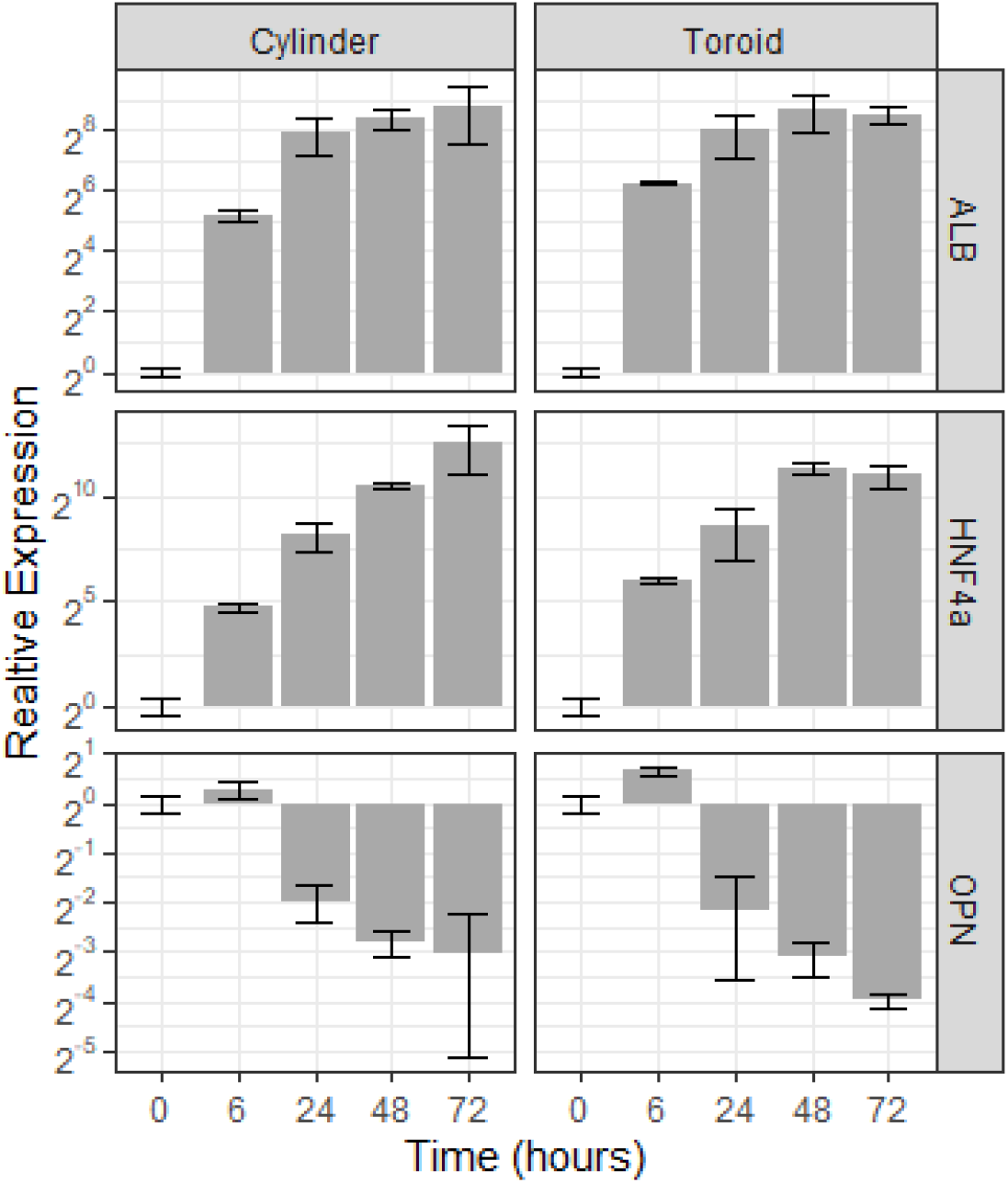
Time course RT-qPCR data for hepatocytic markers Hnf4a albumin (ALB), and osteopontin (OPN). RNA collected from wells containing either only cylinder or only toroid wells. Data normalized against housekeeping gene HPRT expression, and shown relative to expression at time 0, immediately after cells seeded into wells. Error bars shown standard error of the mean.

**Supplementary Table S1.**
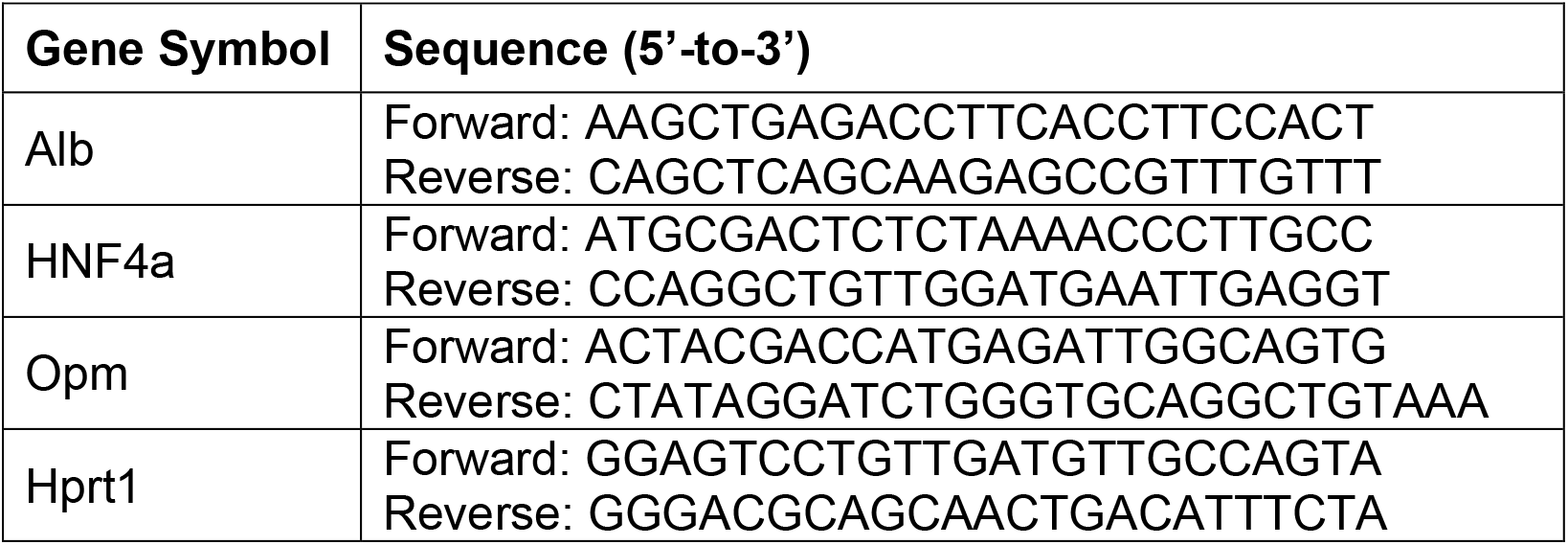
PCR primer pairs

**Supplementary Figure S3.**
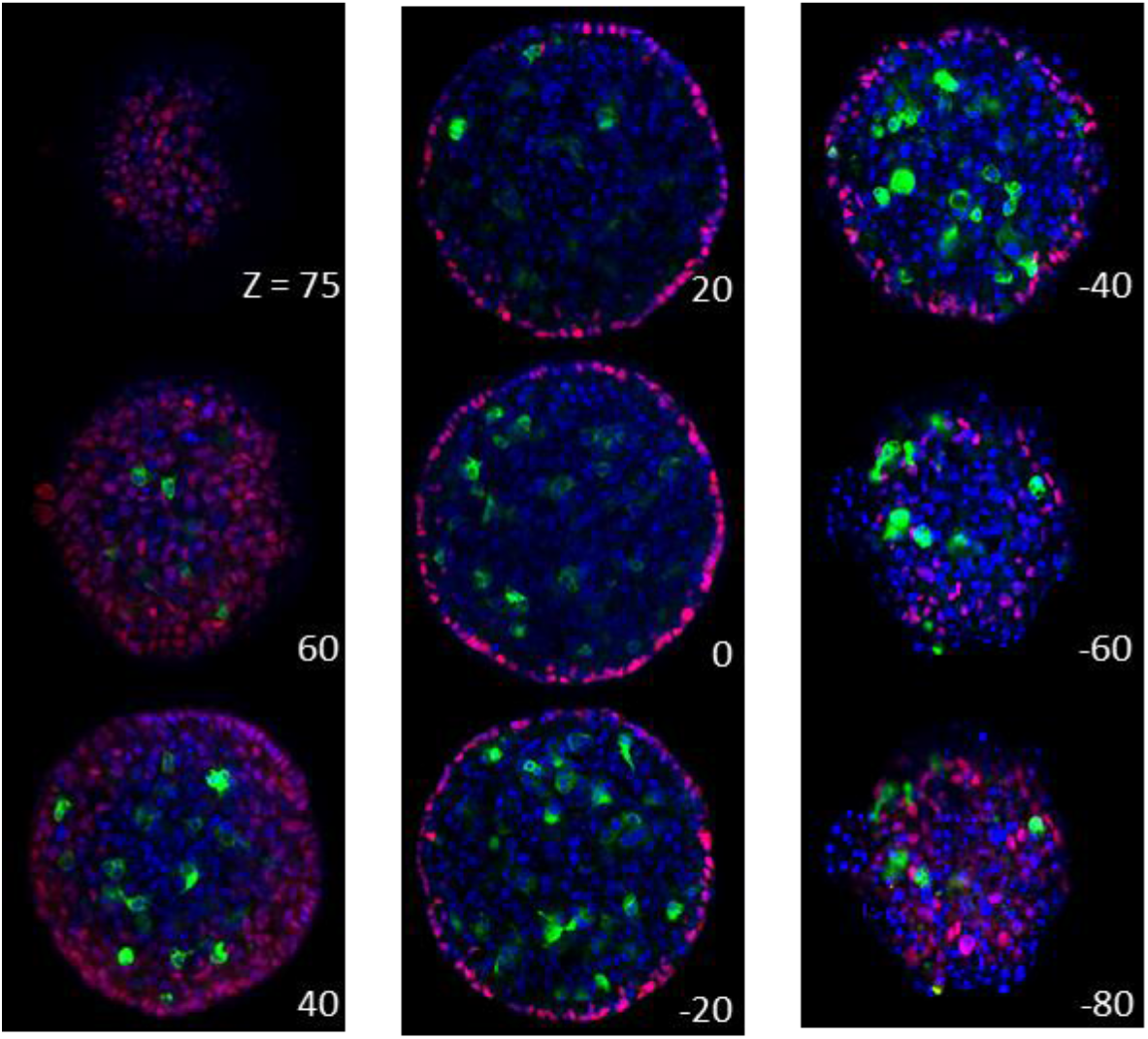
HNF4a and OPN distribution across cylinder microtissue. XY confocal slices of a representative cylinder tissue stained for hepatocytic marker HNF4a (red), biliary marker OPN (green), and with DAPI (blue). Slices labeled by Z distance from the approximate center.

**Supplementary Figure S4.**
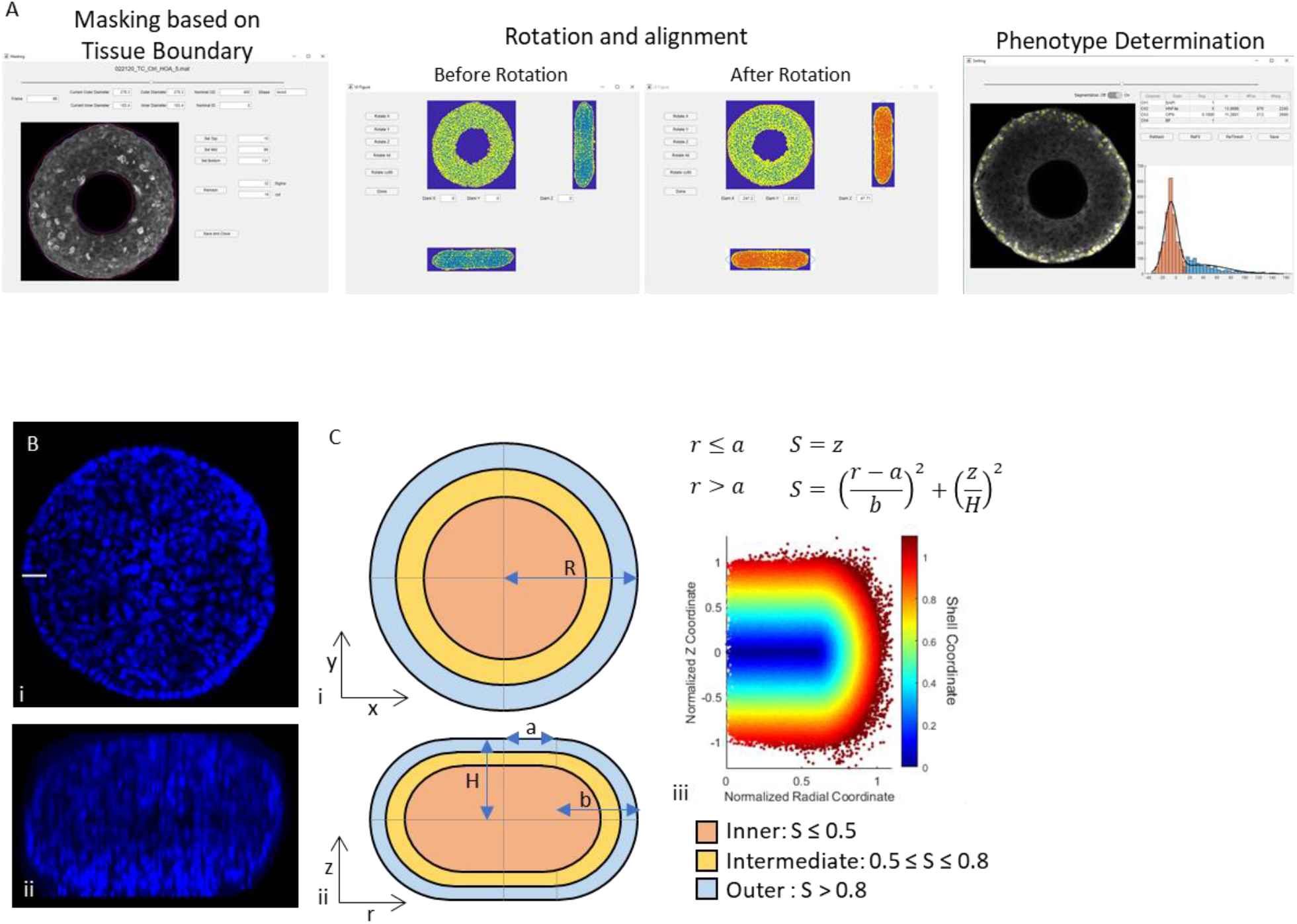
Analysis of segmented data in 3D. A) Outline of process used to identify microtissue boundary, rotate locations in 3D for alignment and normalizing locations for comparison across microtissues, and identifying phenotype based on stain intensity. B) Representative central XY (i) and central XZ (ii) planes of a cylinder tissue stained for DAPI, imaged confocally. C) Approximated geometry of cylinder microtissues, divided into regions in the (i) XY and (ii) ZR planes. iii) Shell coordinate for nuclei plotted in the RZ plane.

**Supplementary Figure S5.**
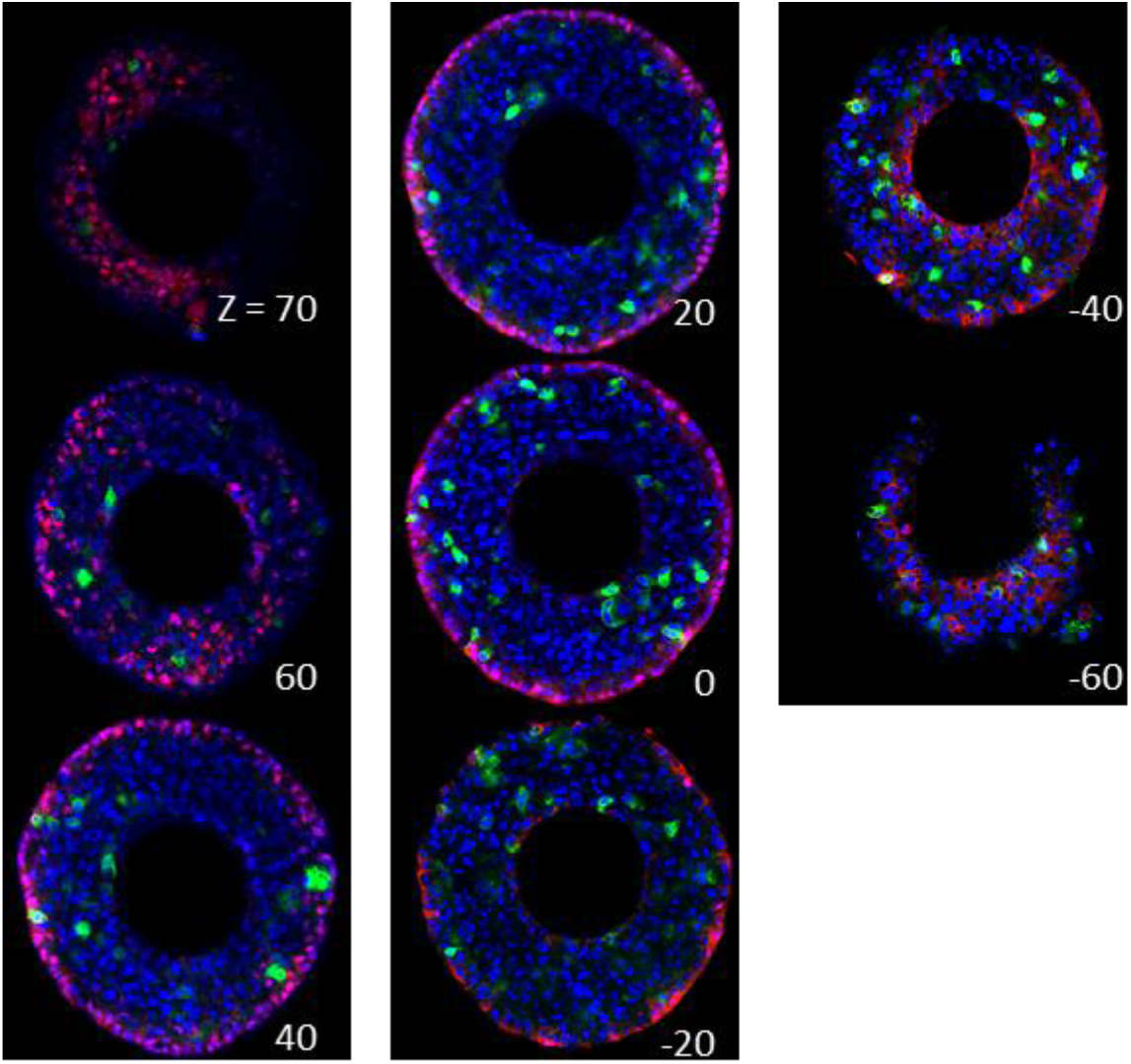
HNF4a and OPN distribution across toroid microtissue. XY confocal slices of a representative toroid microtissue stained for hepatocytic marker HNF4a (red), biliary marker OPN (green), and with DAPI (blue). Slices labeled by Z distance from the approximate center.

**Supplementary Figure S6.**
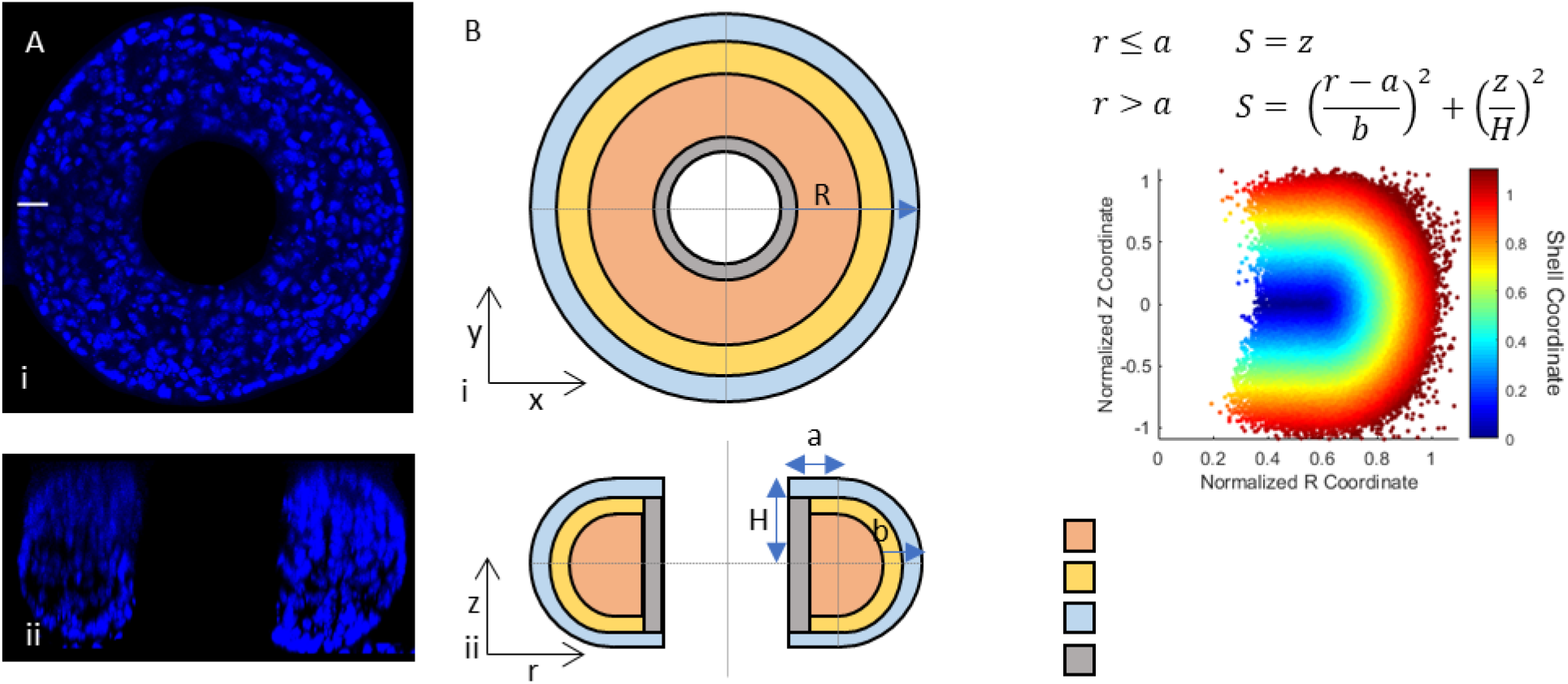
Geometry of toroid microtissues. A) Representative central XY (i) and central XZ (ii) planes of a cylinder microtissue stained for DAPI, imaged confocally. B) Approximated geometry of cylinder microtissues, divided into regions in the (i) XY and (ii) ZR planes. iii) Shell coordinate for nuclei plotted in the RZ plane.

**Supplementary Figure S7.**
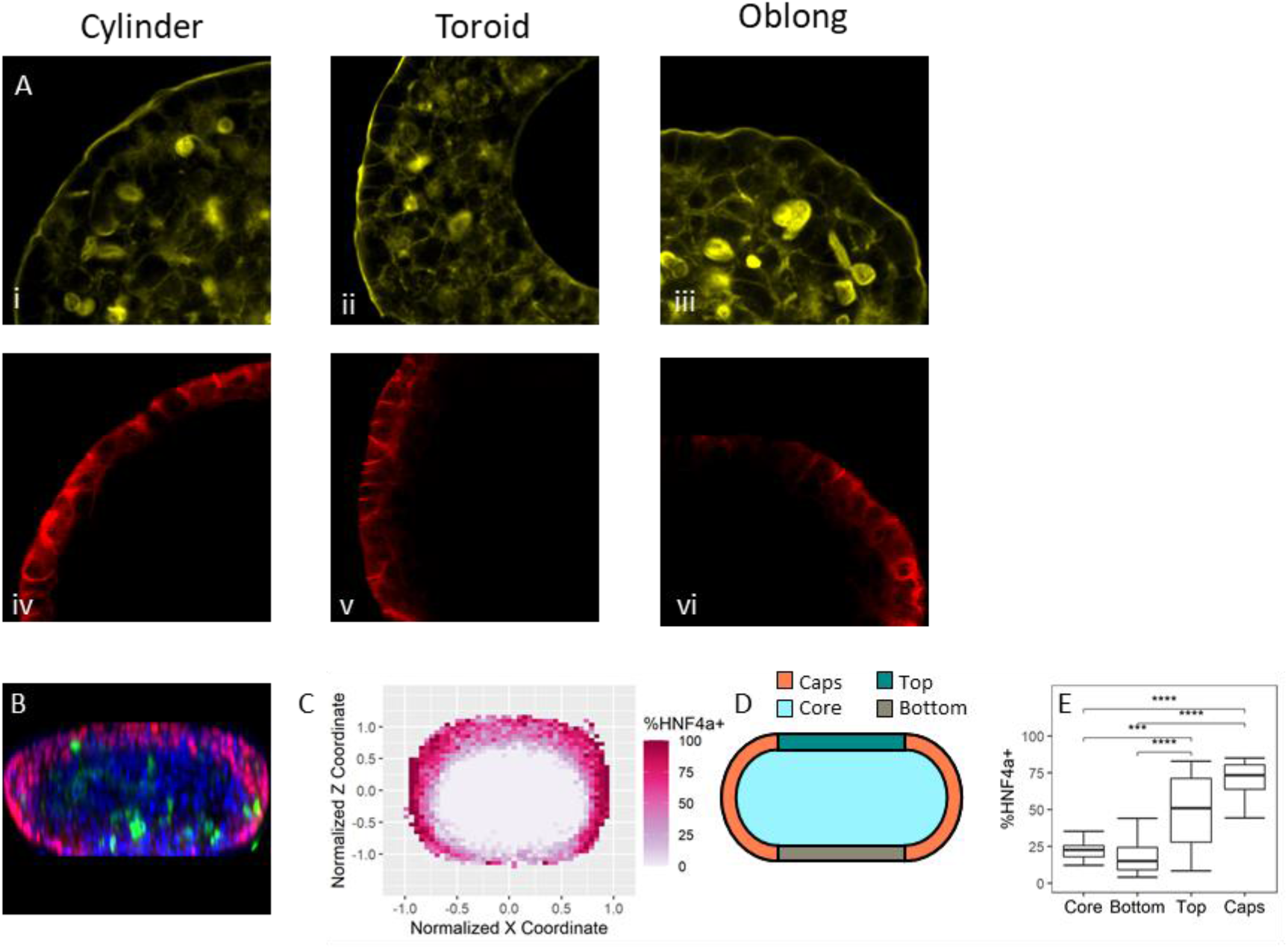
Actin, E-Cadherin, and HN4a in 3D BMEL microtissues. A) Increased magnification view of actin staining in cylinder, toroid, and oblong microtissues. Increased magnification view of E-cadherin staining in cylinder, toroid, and oblong microtissues. B) Confocal XZ image slice of a representative 200 μm wide oblong geometry stained for HNF4a (red), OPN (green), and with DAPI (blue). Scale bar 100 μm. C) Percentage of HNF4a positive cells in XZ space in the center 25% of the microtissue thickness from N=## 200 μm wide toroids. D) Descriptive regions of oblong microtissues. E) Percentage of HNF4a positive cells in the core, flat, and regions of 200 μm geometries. ***p<=0.001, **** p <= 0.0001.

**Supplementary Figure S8.**
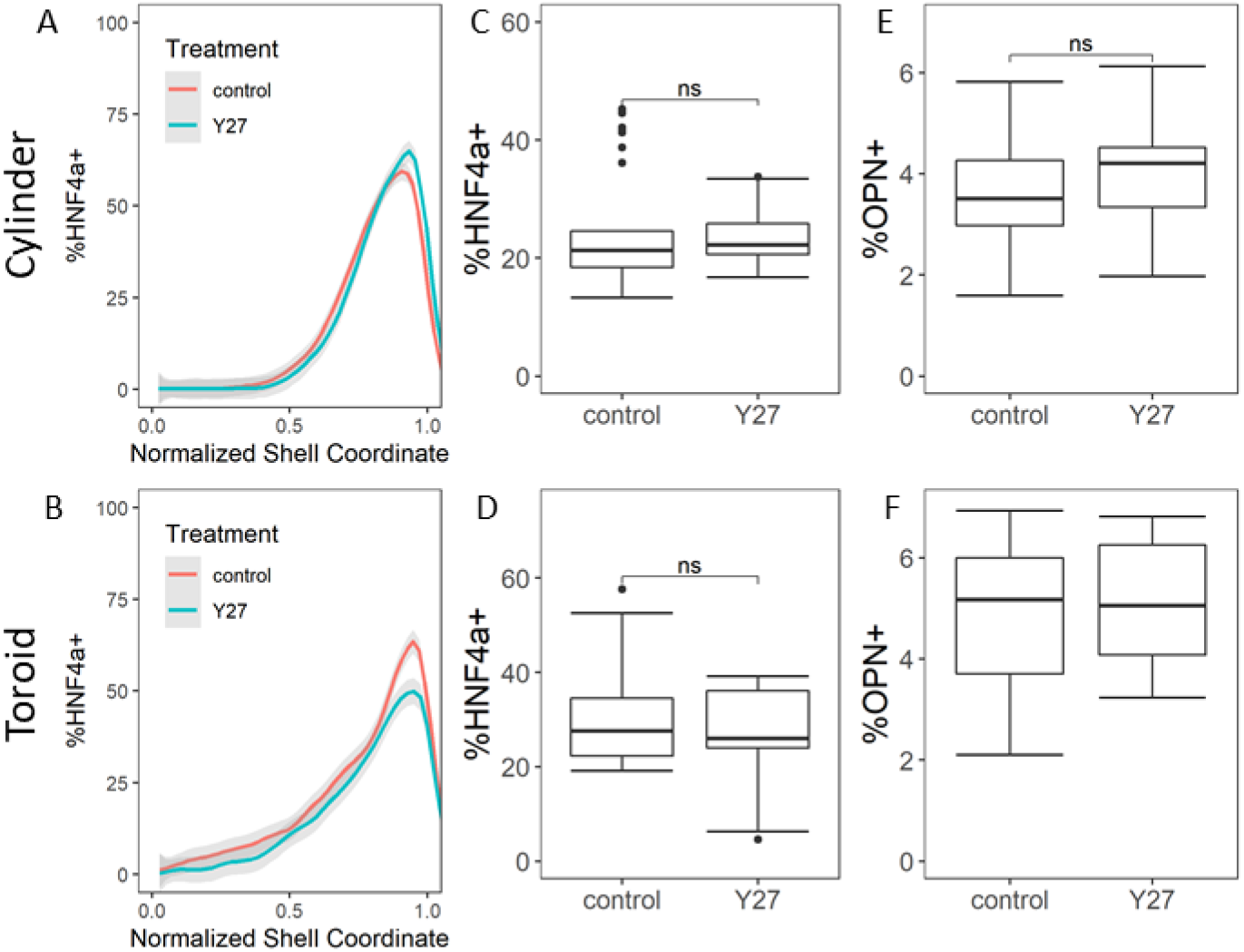
Effects of Y-27632 Treatment. A,B) %HNF4a positive versus shell coordinate for untreated and Y27 treated cylinder and toroid microtissues. C-F) Overall %HNF4a and %OPN positive cells per microtissues in untreated and Y27-treated cylinders and toroids.

**Supplementary Figure S9.**
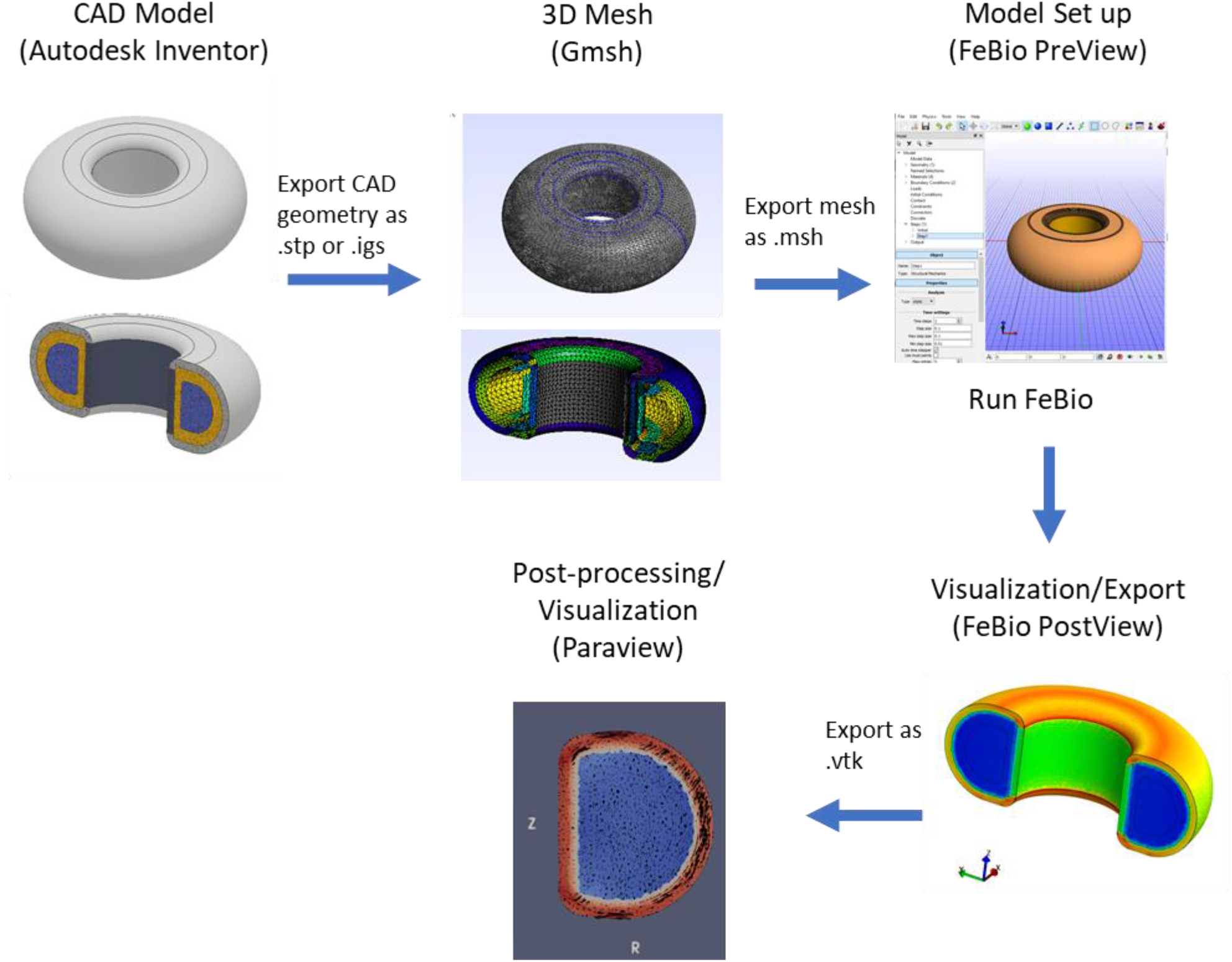
FEM modeling with open-source software. Outline of process and software used to create, run, and analyze FEM models. 3D Tissue geometries were drawn in Autodesk inventor. These geometries were exported as STEP or IGES files. The finite element mesh generator software Gmsh was used to generate mesh files for each geometry. The model was produced using the FEBio software suite. The PreView software was used to import the geometry, set boundary conditions, and specify material properties. The simulation was run using the FeBio solver. Simulation results were visualized in the PostView program, which was also used to export results as a VTK file. Paraview was used to calculate principle stresses, visualize data, and generate figures.

**Supplementary Figure S10.**
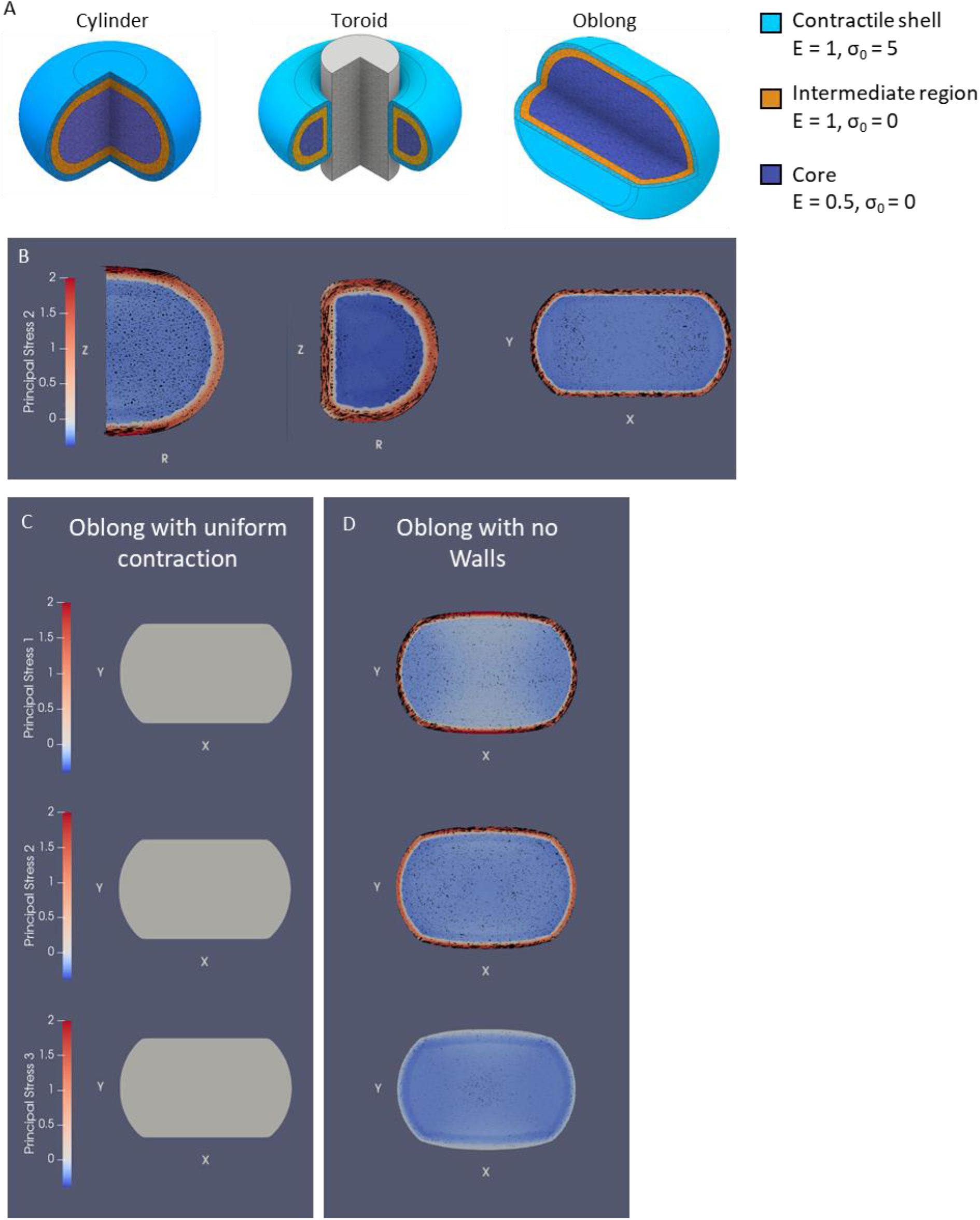
FEM modeling of microtissue contraction. A) 3D models of the cylinder, toroid, and oblong geometries used in the FEM model. Cutaways show the contractile shell, intermediate, and core regions. B) Simulated second principal for an RZ slice of a cylinder and toroid microtissue, and the center XY slice of an oblong geometry. C) Simulated principal stresses for the central XY slice (oblong) with uniform contraction. D) Simulated principal stresses for the central XY slice (oblong) with a passive force, not constrained by walls. Stresses are given in relative units.

## Notes

### Competing Interest Statement

The authors have declared no competing interest.

